# Spacer Extension Reconciles Specificity and Activity in High-Fidelity Cas9 Genome Editing

**DOI:** 10.1101/2025.01.16.633458

**Authors:** Zhike Lu, Rongwei Wei, Rong Zheng, Young-Cheul Shin, Qingfeng Zhang, Jianbo Li, Xiaoqi Wang, Yi Wei, Botao Liu, Yang Chen, Heng Zhang, Hui Chen, Lijia Ma

**Affiliations:** Westlake Laboratory, Hangzhou, Zhejiang, 310024, China; School of Life Sciences, Westlake University, Hangzhou, Zhejiang, 310030, China; Westlake Genetech. LTD., Hangzhou, Zhejiang, 310024, China; Department of Chemical Biology, School of Life Sciences, Southern University of Science and Technology, Shenzhen, Guandong, 518055, China

## Abstract

Engineering high-fidelity CRISPR enzymes often leads to reduced cleavage activity, creating a significant hurdle in balancing nuclease specificity and efficiency for clinical applications. Here, we demonstrate that extending the spacer to 21 or 22 nucleotides restores the impaired cleavage activity of SuperFi-Cas9, an engineered high-fidelity Cas9 variant with seven mutations at the PAM-distal region. Structural and mutational analyses reveal that the spacer extension strengthens additional interactions at the PAM-distal end, stabilizing the nuclease– sgRNA–DNA complex, which appears to be disturbed due to the seven mutations. This approach not only provides a high-fidelity Cas9 with uncompromised efficiency but also introduces a novel strategy to enhance CRISPR complex stability. Our findings offer a promising avenue for precise and efficient genome editing, crucial for advancing CRISPR technologies toward clinical translation.

## Introduction

CRISPR genome editing brings big potential to biomedicine and translational research, as a programmable RNA sequence could guide CRISPR nuclease to edit specific targets simply by sequence complementary in mammalian cells^1,2^. CRISPR nuclease originates from the bacteria’s adaptive immunity for pathogen recognition, which would benefit from a certain level of flexibility on mismatched sequences, but repurposing the nuclease for genome editing requires precise targeting and editing.

In the past decades, structure-guided rationale design aided the engineering of CRISPR nuclease, especially Streptococcus pyogenes Cas9 (Cas9), to eliminate unwanted edits. These rationale designs proposed to generate high-fidelity Cas9 nuclease by mutating residues interacting with either the duplex of sgRNA and targeting strand or the non-targeting strand. The optimal energy thresholds for the Cas9-sgRNA complex to recognize the intended target, for the rehybridization of target and non-target strands, or to active the conformational change of the HNH domain, were discussed and applied to design high-fidelity SpCas9, including SpCas9-HF1^3^, eSpCas9^4^, HypaCas9^5^ et al. Recently, a study using kinetic-guided cryo-EM captured intermediate snapshots when SpCas9/sgRNA binds to mismatched targets, pinpointing a disordered loop in the RuvC domain that specifically contributes to stabilizing a distorted sgRNA-TS duplex^6^. Based on this structural insight, a high-fidelity Cas9, SuperFi-Cas9, was proposed by replacing seven interacting residues in that loop with negatively charged aspartic acid. However, high-fidelity SpCas9, including SuperFi-Cas9, commonly suffers from impaired on-target editing efficiency, and it is extremely unclear why mutations designed from structural insights caused unexpected loss of cleavage activity and whether high-fidelity could live with high efficiency in this powerful nuclease.

Besides working on the nuclease, a few groups reported that engineering the 5’ of sgRNA may decrease undesired mutagenesis of SpCas9. For example, a 5’-truncated sgRNA (tru-sgRNA) was reported to improve editing specificity, hypothesizing that the shorter version of sgRNA decreased the excess affinity provided by the full-length duplex between a 20-nt sgRNA and its target strand^7,8^. Nevertheless, a shorter sgRNA apparently is less unique for its genome target than the 20-nt sgRNA and brings extra challenges to sgRNA design. In contrast, adding a 5’-GG cap out of the 20-nt sgRNA (5’-GGX_20_) also increased the cleavage specificity of Cas9^9^. The additional two guanines facilitate the sequence requirement for U6 or T7 transcription and also lengthen the required complementary region. However, the requirement for genome target complement to 5’-GGX_20_ sgRNA limited its wide application. The hg-sgRNA with designed 5’ stem-loop (hg-sgRNA) also increases editing specificity by modulating R-loop formation^10^, but it is unclear how well the modulation could discriminate on-target and off-target binding.

Here, we propose that high-fidelity SpCas9 variants suffer from reduced cleavage activity as the introduced mutations may change the interactions between the multi-domain enzymes and nucleic acids. Taking SuperFi-Cas9 as an example, we demonstrated that this SpCas9 mutant accommodates a 5’-extended sgRNA better than the canonical 20-nt spacer sgRNA. SuperFi-Cas9 with a 5’-extended sgRNA achieves up to 97.1% cleavage activities of the wild-type Cas9 at endogenous sites and increases 1.87-fold of indel frequencies on average than a 20-nt spacer sgRNA in high-throughput profiling. Structural analysis on SuperFi-Cas9 indicated that the 22-nt spacer sgRNA enhanced interactions between K929, K948, and R951 and the most 5’ end of the sgRNA-TS duplex than the 20-nt spacer sgRNA. Combined with cleavage activities of various SuperFi-Cas9 mutants, we propose that these enhanced interactions improve the stability of the nuclease-sgRNA-TS complex at the PAM-distal region when a 5’-extended sgRNA was used instead of a 20-nt spacer sgRNA. A deep-learning prediction model, AIdit-SuperFi, was built and available at https://crispr-aidit.com/home for 5’-extended sgRNA design. This study brought new insights into the structure-guided SpCas9 design and prompted a high-fidelity nuclease with decent cleavage activity.

## Results

### SuperFi-Cas9 shows a unique nucleotide selectivity to the 21 position at PAM-distal

As the reduction of editing efficiency of SuperFi-Cas9, as well as many other high-fidelity Cas9, is site-dependent, we sought to investigate the target sequence preferences of this nuclease. By using a sgRNA-target paired library, a design allowing high-throughput profiling on cleavage activity of a nuclease from tens of thousands of targets in cells, we quantified the indel frequencies of SuperFi-Cas9 targets cleaved by 91,603 sgRNAs in K562 cells (SuperFi-Cas9 profiling library 1, L1) (**sFig. 1A**). The overall indel frequency of SuperFi-Cas9 is indeed lower than the wild-type SpCas9 and other variants^11^ (sFig. 1A), which is consistent with a previous argument about the impaired cleavage activities of SuperFi-Cas9 and other high-fidelity Cas9 in cell line^12^.

**Fig. 1.**
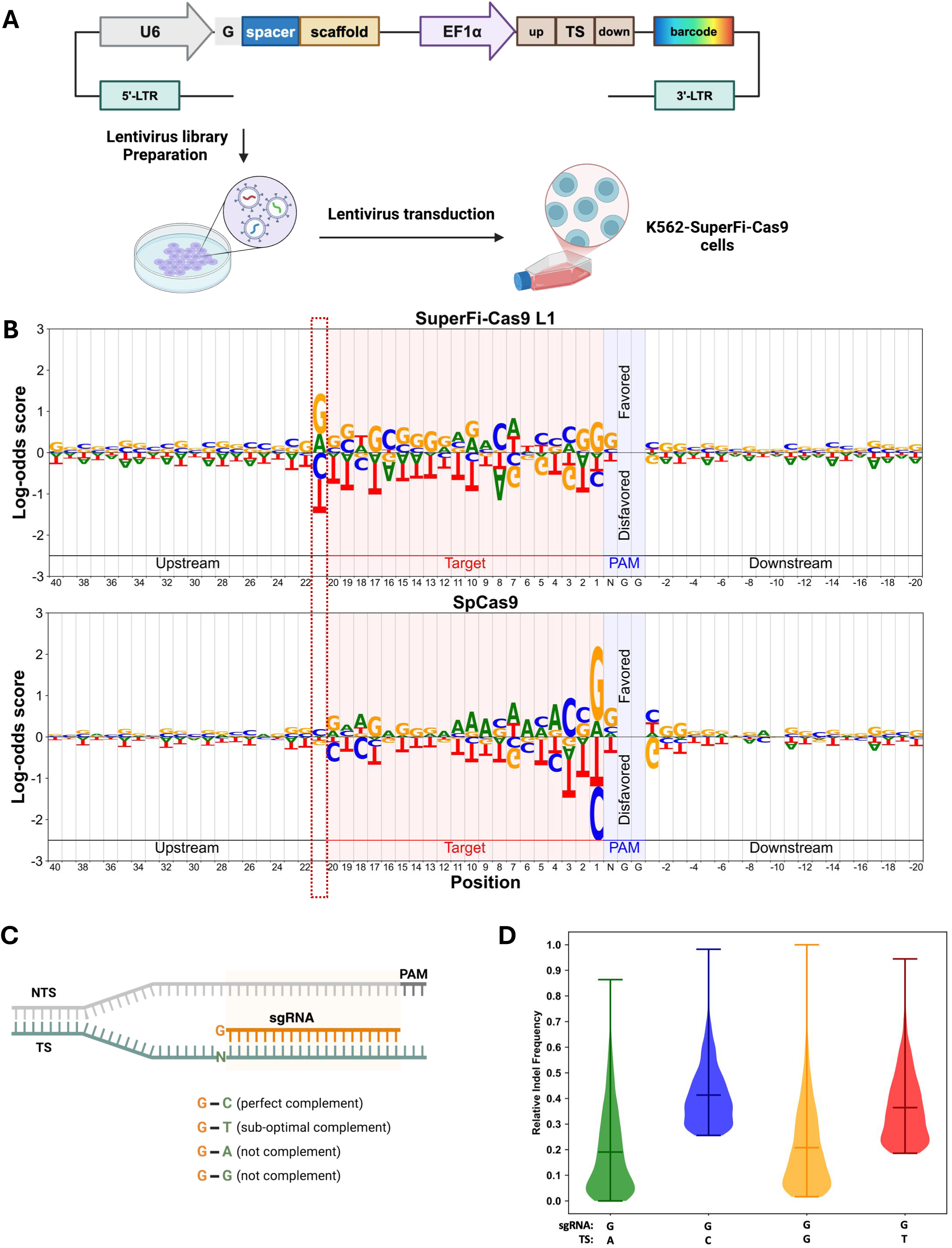

To prompt the use of SuperFi-Cas9, we looked up the target sequence context against their indel efficiencies. By plotting a log-odds score of each nucleotide along the non-target sequence (NTS), we surprisingly found unexpected nucleotide preference at the 21^st^ position, which has been outside of the canonical sgRNA-TS hybridization region (**Fig. 1B**). In contrast, SpCas9 and other variants didn’t show the same nucleotide preference at this position (**Fig. 1B**, **sFig. 1B**). In the high-throughput profiling assay we used, a 5’ leading G was often appended to sgRNA to comply with the sequence requirement of U6 promoter^10,12^ (**Fig. 1C**), however, this appended nucleotide should not influence the cleavage activity of nuclease as it sits outside of the 20-bp sgRNA-TS duplex^10,11,13^. Nevertheless, in SuperFi-Cas9 profiling, when a sgRNA with an additional 5’ leading G targets a 21-bp perfect complementary or sub-optimal complementary sequence, the indel frequencies are significantly higher than that of not complementary target (**Fig. 1D**). Thus, we hypothesized that SuperFi-Cas9 prefers to a sgRNA-TS duplex longer than 20-bp, and its parental nuclease, SpCas9, appears not having this property (**sFig. 1C**).

### Spacer extension increases the cleavage activities of SuperFi-Cas9 at endogenous genomic locations

To investigate whether longer sgRNA-TS duplex could increase the cleavage activities of SuperFi-Cas9, we chose ten endogenous loci in Cas9-expressing K562 cells (K562-SuperFi-Cas9 cells) and designed sgRNAs with different lengths to target them (**Fig. 2A**). In the sgRNA expression cassette, we put a tRNA between the U6 promoter and the spacer of sgRNA, which allows generating extended sgRNAs with designed 5’ sequences that match the target sequences. For each targeted locus, we compared the indel frequencies cleaved by SuperFi-Cas9 with sgRNAs ranging from 20-24 nt, which were named 20-sgRNA, 21-sgRNA, 22-sgRNA, 23-sgRNA, and 24-sgRNA, respectively (s**Fig. 2**).

**Fig. 2.**
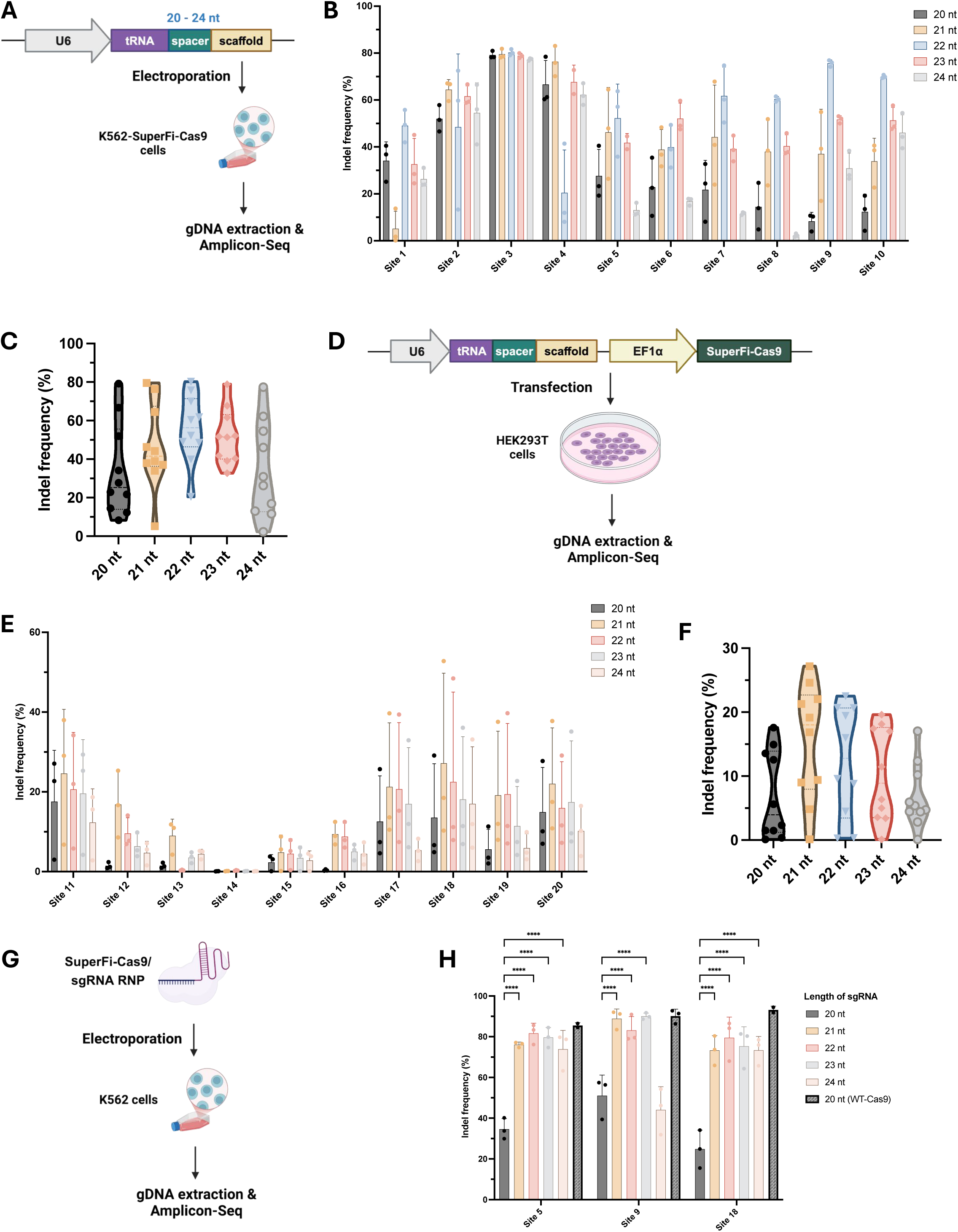

Across all ten targeted loci, the 5’-extended sgRNAs achieved higher indel frequencies compared to their corresponding 20-sgRNA (**Fig. 2B**). The 22-sgRNA presented the highest indel frequencies compared to the other lengths, with mean indel frequencies increased from 33.94% of 20-sgRNA to 55.82% of 22-sgRNA (**Fig. 2C**). Both 21-sgRNA and 23-sgRNA showed increased indel frequencies, while further extending sgRNA to 24-nt appears to have no obvious benefit. To confirm the results, we examined an additional ten endogenous genomic loci in HEK293T cells (**Fig. 2D**). Similarly, the 21-, 22-, and 23-sgRNAs showed higher indel frequencies, and the 21-sgRNA performed the best and achieved 15.46% editing efficiencies compared to 7.01% of the regular 20-sgRNA (**Fig. 2E-F**). Together, 5’-extended sgRNAs effectively increased the editing efficiencies of SuperFi-Cas9 at endogenous sites.

To ensure that the spacer of sgRNA extended to the exact length as we expected, we electroporated K562 cells using ribonucleoprotein (RNP) complexed by synthesized sgRNAs and SuperFi-Cas9 proteins (**Fig. 2G**). At the three loci we examined, the 5’-extended sgRNAs significantly increased the indel frequencies compared to the 20-sgRNA except for the 24-sgRNA at Site 9 (**Fig. 2H**). Compared to the wild-type SpCas9, extending sgRNAs to 21-, 22-, or 23-nt has recovered the editing efficiencies of SuperFi-Cas9 close to the SpCas9/20-sgRNA at Site 5 and Site 9 and about 80% at Site 18. The RNP electroporation data further confirmed that extending the length of sgRNA improves the editing activity of SuperFi-Cas9. At the three sites we tested, SuperFi-Cas9 RNP complexed with extended sgRNA reached comparable indel frequencies of SpCas9/20-sgRNA.

### SuperFi-Cas9 retains high fidelity when utilizing 5’-extended sgRNA

Next, we examined whether SuperFi-Cas9 retains its high fidelity when using 5’-extended sgRNA. We synthesized sgRNAs with one-and two-base mismatches to target Site 5 and Site 9 and examined their cleavage activities in the form of RNP in K562 cells (**Fig. 2G**). At the PAM-proximal region, one-base mismatch sharply reduced the indel frequency of SuperFi-Cas9 (**Fig. 3A-B**). Although the PAM-proximal region is also less tolerant to mismatches by SpCas9, the extent of activity reduction caused by single base mismatch is much more drastic in SuperFi-Cas9^11,13^. At the PAM-distal region, SuperFi-Cas9 is also sensitive to one-base mismatch. Overall, when one mismatch presents within the canonical 20-nt spacer, only two positions in Site 5 and three positions in Site 9 could retain more than half of cleavage activities (**Fig. 3A-B**). Interestingly, a single mismatch at the 19^th^ position of the sgRNA-TS heteroduplex, where SpCas9 and other high-fidelity Cas9 could well tolerate, completely abolished the cleavage activity of SuperFi-Cas9 at both Site 5 and Site 9. In the sgRNA-TS duplex beyond the canonical 20 bp, a 1-bp mismatch at the 21^st^ or 22^nd^ nucleotides seems tolerated by SuperFi-Cas9, but 2-bp mismatches negatively impacted the indel frequencies. At Site 5, CA to TG mismatches were better tolerated than CA to GT mismatches, presumably because the mismatched G at the 21^st^ position of the spacer provides a non-canonical base pairing with the T on the target DNA strand. Together, the 5’-extended sgRNA retains the high fidelity of SuperFi-Cas9 with a unique stringency at the PAM-distal position 19.

**Fig. 3.**
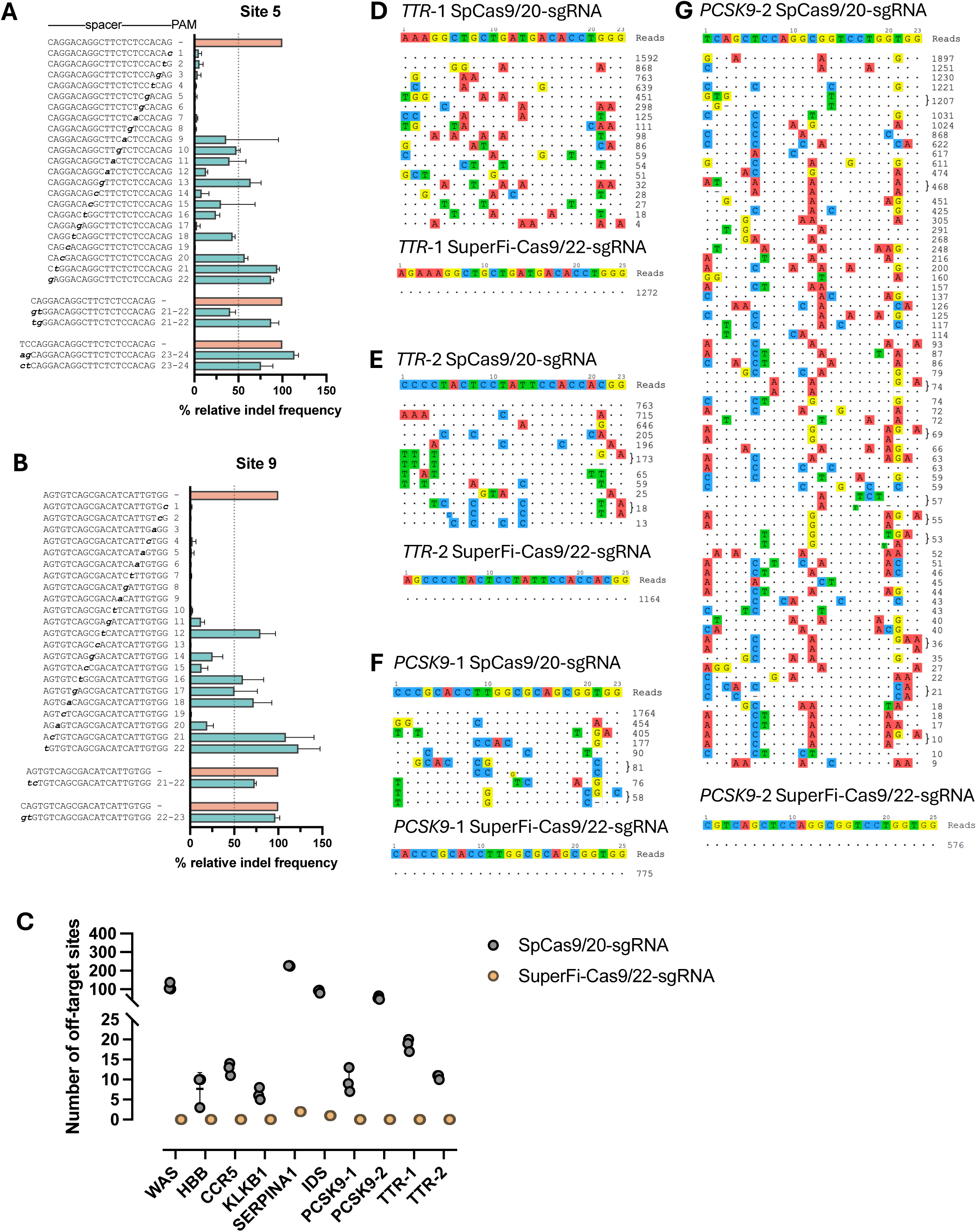

To assess the global off-target editing of SuperFi-Cas9, we conducted GUIDE-seq on genes used in CRISPR therapy. We chose ten sgRNAs targeting eight genes, including WAS, HBB^14^, CCR5^15^, PCSK9^16^, TTR^17^, KLKB1^18^, SERPINA1^19^, and IDS^20^. When using SuperFi-Cas9 as nuclease, GUIDE-seq detected no off-target from eight sgRNAs, one off-target from the 22-nt sgRNA targeting IDS, and two off-targets from the 22-nt sgRNA targeting SERPINA1 (**Fig. 3C**, **Supplementary Table 1)**. In contrast, SpCas9 generates off-targeting edits from all ten 20-nt sgRNAs, with 6 to 226 off-targets per target. The sgRNA TTR-1 is selected from NTLA-2001^17^, which is currently in clinical trial to develop medicine to treat transthyretin amyloidosis by systemic administrating lipid nanoparticle (LNP) encapsulating SpCas9/20-sgRNA ribonucleoprotein (RNP). NTLA-2001 reported 12 GUIDE-seq off-targets, and our GUIDE-seq experiment detected 18. However, no off-targeting edits were detected using SuperFi-Cas9 and a 22-sgRNA (**Fig. 3D**). An alternative sgRNA TTR-2 was designed to target the same gene, and GUIDE-seq revealed similar results of the SpCas9/20-sgRNA and SuperFi-Cas9/22-sgRNA (**Fig. 3E**). Similarly, sgRNA PCSK9-1 is selected from VERVE-101 and VERVE-102, which are clinical trials for investigating treatments of familial hypercholesterolemia. The sgRNA PCSK9-1 is used with ABE8.8 base editor mRNA in an LNP formulation. ONE-seq reported 45 off-targets of SpCas9/20-sgRNA and our GUIDE-seq detected 10. We detected no off-targets using SuperFi-Cas9/22-sgRNA (**Fig. 3F**). GUIDE-seq on an alternative PCSK9 targeting (Fig. 3G) and other six targets revealed the same pattern (s**Fig. 3**). The very few numbers of GUIDE-seq off-targets highlight the high fidelity of SuperFi-Cas9 with a 5’-extended sgRNA on CRISPR therapy targets.

### High-throughput profiling confirmed the restored SuperFi-Cas9 cleavage activities by spacer extension

By spacer extension, we have shown that SuperFi-Cas9 restored cleavage activity at selected endogenous sites. To analyze the impact of spacer extension to the nuclease in high-throughput, we generated a SuperFi-Cas9 profiling library containing sgRNAs with 19-24 nt spacer transcribed from a U6-tRNA-sgRNA architecture (SuperFi-Cas9 profiling library 2, L2) (**Fig. 4A**). We compared the indel frequencies of sgRNAs in different lengths on 3,632 target sites, at which SpCas9 can achieve >50% indel frequencies^11^. Consistently with the observations at the twenty endogenous targets, 21-sgRNA and 22-sgRNA significantly improved the indel frequencies of SuperFi-Cas9 compared to other lengths in high-throughput profiling (**Fig. 4B**). To exclude the potential interference or bias caused by the tRNA architecture, we constructed an independent profiling library, in which different lengths of sgRNAs were directly transcribed from a U6 promoter without tRNA cleavage (SuperFi-Cas9 profiling library 3, L3) (**Fig. 4C**). The 21-sgRNA and the 22-sgRNA from the L3 are the best-performed categories among the other lengths, which again confirmed that spacer extension could restore the indel frequencies of SuperFi-Cas9.

**Fig. 4.**
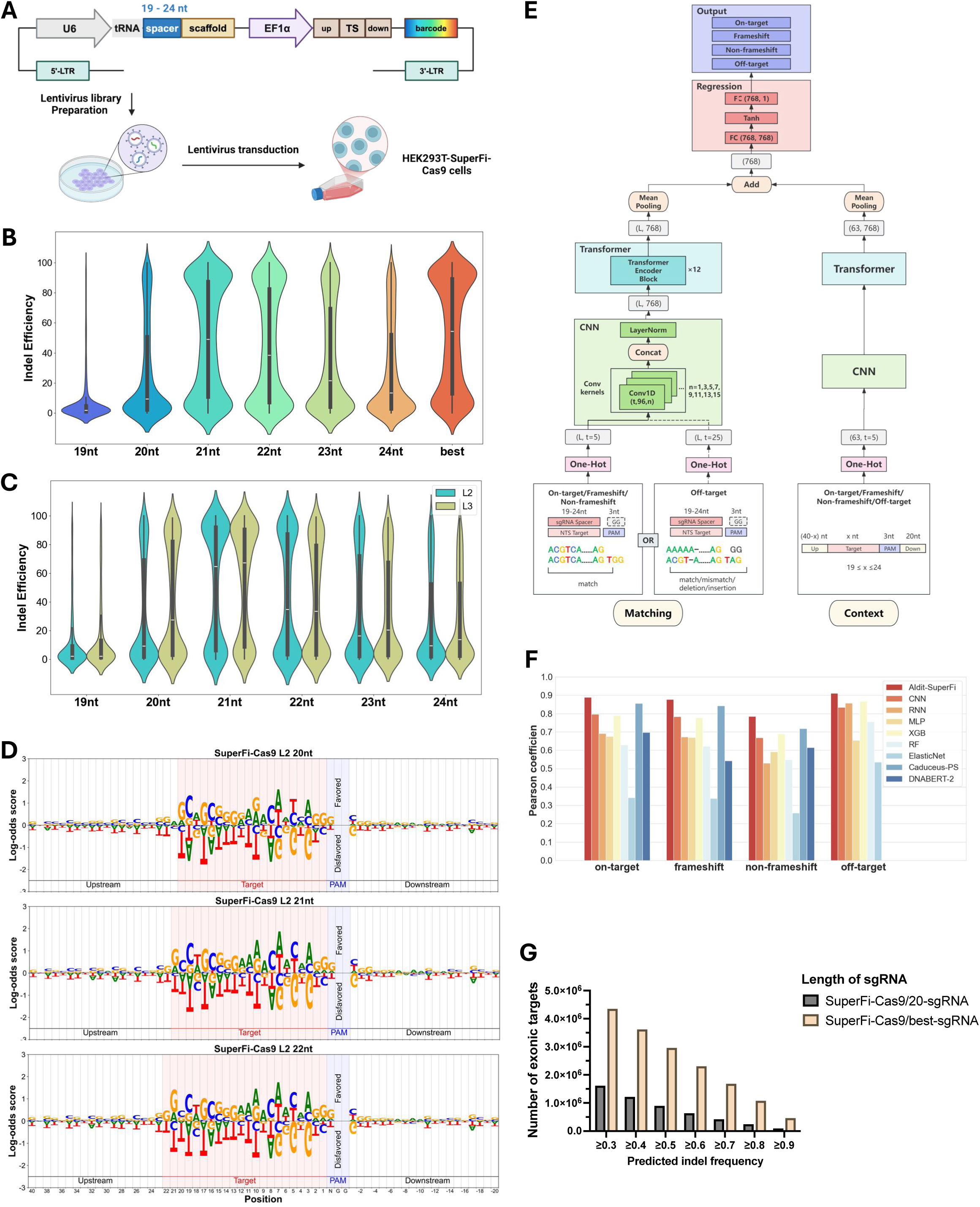

Noticeably, the indel frequencies of sgRNAs showed a binary distribution, indicating that the cleavage activities of SuperFi-Cas9 could be restored by spacer extension at some targets while remaining low at others. By plotting the ratio of nucleotide in high-activity and low-activity sgRNAs, we found that SuperFi-Cas9 generally prefers sgRNAs with cytosine and guanine at the PAM-distal region (**Fig. 4D**, **sFig. 4-5**), while SpCas9 and other variants prefer adenosine and guanine (**Fig. 1B**, **sFig. 1B**). Indeed, the number of CG pairs at the PAM-distal region of SuperFi-Cas9 is positively correlated with its indel frequency, while no such correlation was seen in SpCas9 (**sFig. 6-7**). Similarly, the free energy of the sgRNA-TS duplex predicted by the nearest-neighbor model^21^, which reflects its stability, also shows a positive correlation with the indel frequencies of sgRNA of SuperFi-Cas9 but not SpCas9 (**sFig. 8-9**). This nucleotide preference and correlation with indel frequencies indicate that SuperFi-Cas9 appears to require a more stable sgRNA-TS duplex at the PAM-distal region.

**Fig. 5.**
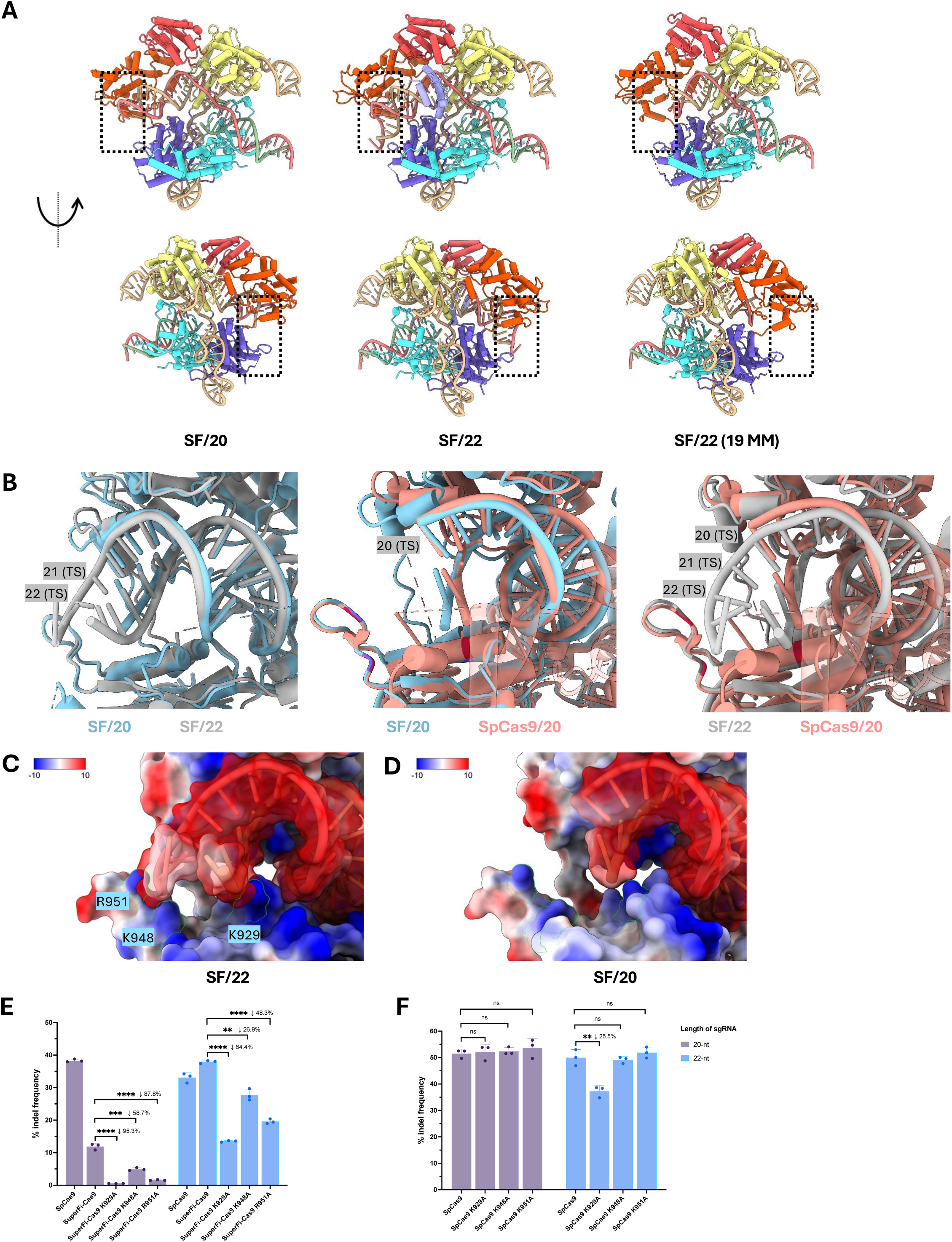

To aid the design of the best-performed sgRNA for SuperFi-Cas9 among different spacer lengths, we trained AIdit-SuperFi, a deep-learning (DL) model to predict cleavage activities and editing specificity for sgRNA of SuperFi-Cas9. The editing specificity prediction model was trained from an additional off-target SuperFi-Cas9 library (L4) we generated for this study. AIdit-SuperFi is based on Convolutional Neural Networks (CNN) and Transformer Encoder^22^ and makes the prediction based on sgRNA, PAM, target, and flanking genomic sequences (**Fig. 4E**, **sFig. 10A**). The model predicts the indel frequency, frameshift, non-frameshift, and off-target efficiency from the user-defined locus. On all four tasks, AIdit-SuperFi consistently outperforms the other baseline models (**Fig. 4F**, **Supplementary Table 2)**. Moreover, on all off-target prediction sub-tasks, the AIdit-SuperFi model also performs best (**sFig. 11**, **Supplementary Table 3)**. The prediction model is available at a user-friendly GUI at https://crispr-aidit.com/home, which could help with designing the best spacer length to conduct high-efficiency and high-fidelity CRISPR gene editing by SuperFi-Cas9. AIdit-SuperFi model predicts a sufficient amount of 5’-extended sgRNA that could produce decent indel frequencies in exons of human protein-coding genes (Fig. 4G).

### Cryo-EM revealed enhanced interactions between SuperFi-Cas9 and 22-sgRNA-TS duplex

Curiously, we sought to understand why the 5’-extended sgRNA improves the cleavage activity of SuperFi-Cas9 compared to using a canonical 20-nt spacer. We resolved the structures of SuperFi-Cas9 complexed with a 20-or 22-nt sgRNA and their matched dsDNA targets (**Fig. 5A**). We also resolved the structure of SuperFi-Cas9 complexed with a 22-nt sgRNA with one mismatch at the 19^th^ position, considering the dramatic activity loss caused by this mismatch (**Fig. 3A-B**). Cryo-EM reconstructions of SuperFi-Cas9 with 20-nt, 22-nt, and 22-nt mismatched sgRNA (3.5 Å, 3.6 Å, and 3.3 Å respectively) showed similar domain architectures (**sFig. 12-14**). Compared to SuperFi-Cas9/20-sgRNA (SF/20), the sgRNA-TS duplex of SuperFi-Cas9/22-sgRNA (SF/22) extended further at the PAM-distal region. When the mismatch was presented at the 19^th^ position, the PAM-distal heteroduplex was disordered, as shown in the structure of “SF/22 (19 MM)” (**Fig. 5A**, **sFig. 15**). Except for the PAM-distal heteroduplex, the three structures are overall similar, and the HNH domain is disordered in most subclasses (**sFig. 16**).

The structural alignments showed that the extended 21^st^ and 22^nd^ sgRNA-TS base pairs of SF/22 came near residues M939 to K954, a loop in the RuvC domain (**Fig. 5B**, **sFig. 17**). In that loop, K948 and R951 are two positively charged residues that are located most closely to the extended 21^st^ and 22^nd^ of the negatively charged sgRNA-TS duplex and the unpaired 23^rd^ and 24^th^ nucleotides at TS strand (**Fig. 5B-C**). Both structural alignment and the Coulombic electrostatic surface potential map indicated an enhanced electrostatic interaction between the 5’ of sgRNA and K929 in the structure of SF/22 compared to SF/20 (**Fig. 5B-D**). These three residues (K929, K948, and R951), although not in contact with nucleic acids in our experimental structure, suggested different interaction strengths at the PAM-distal region between SF/20 and SF/22, which motivates us to further explore their influences on the editing efficiencies of SuperFi-Cas9.

Indeed, the K929A mutant significantly decreased the cleavage activity of SuperFi-Cas9/22-sgRNA by 64.4%. Mutations at K948 and R951, the two residues in the loop approached by the extended sgRNA-TS duplex, also caused cleavage activity loss but to a less extent (26.9% and 48.3%, respectively) (**Fig. 5E**). In contrast, none of these mutants changed the cleavage activities of SpCas9/20-sgRNA (**Fig. 5F**). These results aligned with the observation from the structural alignment that spacer extension enhanced the interactions between K929 and SuperFi-Cas9/22-sgRNA, which potentially stabilized the PAM-distal region and made K929A an essential mutation to the cleavage activity of SuperFi-Cas9/22-sgRNA. The potential electrostatic interactions between K948/R951 and SuperFi-Cas9 are likely not critical to SpCas9/20-sgRNA.

To further elucidate whether the activity losses with K929A, K948A, and R951A are unique to SuperFi-Cas9 or the spacer extension, we looked up the cleavage activities of SuperFi-Cas9 with a canonical 20-sgRNA and SpCas9 with an extended 22-sgRNA. The dramatic activity losses of SuperFi-Cas9/20-sgRNA suggested a fragile interface at its PAM-distal region, where each of the three mutations seems to greatly impact the cleavage activity (**Fig. 5E**). In contrast, although hydrogen bonds were observed between K929 of SpCas9 and the phosphate backbone of sgRNA at the 19th and 20th positions^23^, only 25.5% activity loss was caused by K929A mutation. No activity change was caused by K948A and R951A of SpCas9/22-sgRNA. Together, K929, K948, and R951 are more essential to the cleavage activities of SuperFi-Cas9 than SpCas9, regardless of using a 20-or 22-nt spacer.

## Discussion

Given the potential of CRISPR technology in treating genetic disorders and improving precision in cancer therapies, developing high-fidelity nuclease variants with retained on-target activity is paramount. In this study, we successfully restored the cleavage activity of SuperFi-Cas9, a rationale-designed high-fidelity Cas9 variant, while retaining its editing specificity. By spacer extension, SuperFi-Cas9 achieves up to 97.1% editing efficiency of SpCas9 in cells while even responding to a single mismatch in both the seed region and the distal end of the duplex. High-throughput profiling shows that spacer extension increases 1.87-fold cleavage activities of a canonical 20-nt spacer on average and reveals a nucleotide preference for cytosine and guanine at the PAM-distal region. Spacer extension also allows the use of SuperFi-Cas9 as base editors (sFig. 18-19), and the editing window of SuperFi-CBE was extended both upstream and downstream of the canonical editing window. Moreover, structural analyses of SuperFi-Cas9 underscore the importance of the sgRNA-TS heteroduplex at the PAM-distal end, where enhanced interactions between the nuclease and nucleic acids were observed for 22-sgRNA. SuperFi-Cas9 mutants with a single mutation at K929, K948, and R951 significantly reduced the cleavage activities of SuperFi-Cas9 but not SpCas9, except for a mild activity loss of SpCas9-K929A/20-sgRNA. Overall, the spacer extension appears to stabilize the structure at the PAM-distal through the enhanced interactions between SuperFi-Cas9 and sgRNA-DNA heteroduplex.

This study provides a novel strategy to increase the cleavage activities of SpCas9 through spacer 5’-extension. Engineering sgRNA can tune the specificity of CRISPR nucleases, but little has been done to tune their cleavage activity, although the advantage of using longer sgRNA is obvious in the context of pursuing high-fidelity. Unlike 5’-GGX_20_ sgRNA^9^, which brings additional sequence requirements to sgRNA design, the use of 21-or 22-nt sgRNA increased the probability of finding unique targets in the genome. For instance, extending sgRNA from 21 to 24-nt turned 5,370 ClinVar mutations into unique CRISPR targets, accounting for 23.5% of non-unique targets left by the 20-nt sgRNA. The ability of SuperFi-Cas9 to efficiently target unique genomic loci enhances the potential for precise genome editing in therapeutic applications, such as correcting genetic disorders.

The spacer extension also provides broad insights into the structure-guided rationale design of high-fidelity SpCas9 variants. In many cases, this type of design refers to the structure snapshot of the product state^3,4^. A common practice is to mutate the residues interacting with either the sgRNA-TS heteroduplex or the non-target strand, with the hope of optimizing the balance of energies provided by the interactions and necessitated by nuclease recognition and cleavage. However, because SpCas9 requires multiple steps of domain rearrangement for effective target recognition and cleavage, the impact of these mutations can be complex. Spacer extension provides an alternative approach to balance the efficiency and specificity by adjusting the interactions between the nuclease and nucleic acids, which has been proven effective in tuning the disturbed interactions at the PAM-distal end in this study.

Introducing seven aspartic acids (7D) into a protruded loop at the PAM-distal end not only disrupts the interactions in the mismatched conformation but also forms electrostatic repulsion between the loop and the PAM-distal heteroduplex. The repulsion likely shifts the properly positioned complex and leads to the loss of structural stability when using the 20-nt spacer, which significantly impairs the cleavage activity of SuperFi-Cas9 in cells. Using the 22-nt spacer strengthens the interactions and compensates the repulsion and, thus, the stability at the PAM-distal region, which is essential to the nuclease’s overall functionality. The PAM-distal region is where the nuclease-sgRNA-TS complex undergoes structural checkpoint and rearrangement during target binding and cleavage. The stabilized complex likely allows the nuclease to maintain a more rigid and active confirmation during cleavage. However, more investigations will be needed to elucidate the full mechanism.

In conclusion, our study offers a practical approach to achieving high editing efficiency using a high-fidelity Cas9 variant while emphasizing the critical role of the extended heteroduplex at the PAM-distal end in stabilizing structural interactions. By elucidating these mechanisms, we contribute to the rational design of Cas9 variants with high editing specificity and pave the way for more effective and precise CRISPR applications in biomedicine and beyond.

## Methods

### Establish sgRNA-target paired profiling library

We used the sgRNA-target paired oligo pool designed by Zhang et al^11^ to construct a lentivirus library and profile indel frequencies of SuperFi-Cas9.

In brief, a barcode oligo with 20 nt random nucleotides was amplified by PCR (primers: Barcode-PCR-F and Barcode-PCR-R), and the purified PCR products were cloned into the backbone vector (ON-OFF-Backbone_lentiGuide-Puro-Mkate2) by Golden Gate. The primers used for the barcode oligo amplification and the sequence of the barcode oligo are listed in **Supplementary Table 4**. The oligo pool was synthesized by a vendor (GenScript) and was used as templates in PCR, and the PCR products were purified and cloned into the barcode vector by Golden Gate. The product vector library was further ligated into the ON-OFF-Insert-Scaffold vector by Golden Gate. Two independent plasmid libraries were prepared, and each library was used in one replicate of the two biological replicates of high throughput screening.

NGS library of the gRNA-target pairs and barcode cassette in the final plasmid library was established and sequenced as a starting reference. The PCR reaction included 50 μL reaction included 40 ng plasmid, 250 nM forward primer (Fwd-libseq), 250 nM reverse primer (Rev-libseq), 25 µl of NEBNext Ultra II Q5 Master Mix, and nuclease-free H_2_O. The PCR program was set as following: 30 s at 98 °C; 6 cycles of 10 s at 98 °C, 30 s at 64 °C, and 10 s at 72 °C; 8 cycles of 10 s at 98 °C, 30 s at 71 °C, and 10 s at 72 °C; 2 mins at 72 °C; 4 °C hold. The PCR products were purified using AMPure XP beads and subjected for NGS sequencing. The primers used for the NGS library preparation are listed in **Supplementary Table 4**.

To prepare the lentivirus library, the transfer plasmid, psPAX2, and pMD2.G were mixed at a weight ratio of 5:3:2, and a total of 192 ug plasmid mixture were transfected to HEK293T cells at 90% confluent in a T175 flask using the calcium phosphate method according to the published protocols^22, 23^. The culture media was changed eight hours after transfection, and the cells were washed with PBS to remove residual plasmids and transfection reagents. After 48 hours and 72 hours of transfection, we collected the supernatants containing the lentivirus. The supernatants were filtered through a 0.45 μm polyvinylidene fluoride filter and spun in an ultracentrifuge at 20,000 rpm for 2 hours. The precipitation was resuspended in 1 mL PBS and stored at -80 °C in the proper size of aliquots.

After we understood that SuperFi-Cas9 prefers a 5’-extended sgRNA, we designed a new sgRNA-target paired oligo pool, in which 19-24 nt sgRNAs were designed to edit one target sequence. The construction of the plasmid library and the lentivirus library were the same as described above.

### High throughput profiling of SuperFi-Cas9/sgRNA in K562 cells and HEK293T cells

To profile the indel frequencies of SuperFi-Cas9 in high throughput, we transduced the lentivirus libraries containing the sgRNA-target paired sequences into the SuperFi-Cas9-expressing K562 cells (K562-SuperFi-Cas9) or HEK293T cells (HEK293T-SuperFi-Cas9) at MOI=5 by spinfection, which was achieved by spinning cells at 600g for 2 hours at 32 °C. At the end of spinfection, 1 mL of pre-warmed RPMI with 15% FBS and 8 μg/mL polybrene was added into each well immediately. 36 hours post-transduction, the transduced K562-SuperFi-Cas9 cells were selected under 3 μg/mL puromycin for 2 days. 84 hours post-transduction, the K562-SuperFi-Cas9 cells or HEK293T-SuperFi-Cas9 that successfully transduced by the lentivirus were harvested and ready for NGS library preparation.

To prepare the NGS library for the gRNA-target pair and barcode cassette, genomic DNA (gDNA) was extracted from the collected cells using DNeasy Blood & Tissue Kits (Qiagen). The gRNA-target pairs and barcode cassette were amplified by multiple PCR reactions to use up the extracted gDNA. Each 50 μL reaction included 2.5 μg gDNA, 250 nM forward primer (Fwd-libseq), 250 nM reverse primer (Rev-libseq), 25 μL NEBNext Ultra II Q5 Master Mix (New England Biolabs), and nuclease-free H_2_O. The PCR program was set as following: 30 s at 98 °C; 10 cycles of 10 s at 98 °C, 30 s at 64 °C, and 10 s at 72 °C; 12 cycles of 10 s at 98 °C, 30 s at 71 °C, and 10 s at 72 °C; 2 mins at 72 °C; 4 °C hold. The PCR products were combined and concentrated to 100-200 μL by Amicon Ultra 0.5-mL Centrifugal Filters (Sigma). SPRI beads (Beckman Coulter) were used to select the products with the expected size at 420 bp. The size-selected products were subjected to NGS sequencing. The primers used for the NGS library preparation are listed in **Supplementary Table 4**.

### Bioinformatic analysis of the high throughput screening data

The data analysis was conducted following the same pipeline as described in Zhang et al^11^. In brief, sequencing reads were subjected to adaptor removal, alignment, and quality control using in-house Python scripts. The gRNA-target and barcode cassette in the plasmid library were used as starting reference to determine the sequence species that pre-existed before the high throughput screening in cells, and thus, non-existed gRNA-target and barcode cassette were considered as errors introduced by plasmid or PCR amplification and removed from the screened (edited) library. The gRNA-target and barcode cassette in the plasmid library was also used to determine the barcode (BC) sequences associated with a gRNA. In this study, we used a cutoff of > 100 sequencing reads and > 5 BC reads to filter high-quality reads. In total 62,083 gRNA passed quality filtration and retained in the following analysis and modeling.

To determine the indel frequency of each gRNA on their synthetic target, we counted the number of sequencing reads aligned to the 20 bp synthetic target sequence in the edited library. The indel frequency of each gRNA was calculated as:

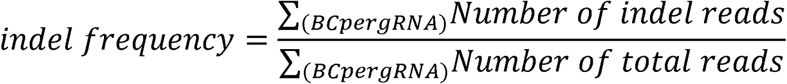

### Measure the editing efficiency at endogenous genomic locations

We randomly chose twenty targets (Site 1 to Site 20) from the sgRNA-target library to quantify the editing efficiencies of sgRNAs at their endogenous sites. For each target, we designed sgRNAs with 21-24 nt matched sequences and a 21 nt gRNA with an unmatched 5’ nucleotide (**Supplementary Table 1**).

To target Site 1 to Site 10, we electroporated the sgRNA-expressing vector into K562-SuperFi-Cas9 cells. A tRNA was placed between the U6 promoter and gRNA, allowing the incorporation of the designed leading nucleotide at the 5’ of the expressed gRNA. K562-SuperFi-Cas9 cells were cultured in 10% FBS and 5 μg/mL blasticidin in RPMI1640 (Sigma R8758). To conduct electroporation, 1 million cells were resuspended in 98 μL LONZA SF Nucleofector Solution (with the provided supplement supplied in the SF Cell Line 4D-Nucleofector™ X Kit) together with 5 μg gRNA plasmids (V6-opt-tRNA-gRNA). Cells were electroporated using program FF-120 of the LONZA 4D-Nucleofector™ X Unit following the manufacturer’s instruction. Pre-warmed (37 °C) culture media was added to cells immediately after electroporation, and cells were transferred and cultured in a 24-well plate. Cells were harvested 48 hours post-electroporation.

To target Site 11 to Site 20, we transfected an all-in-one vector (SuperFi-Cas9-tRNA-gRNA-all-in-one) to HEK293T cells. HEK293T cells were seeded into a 24-well plate at the cell density of 1.5 × 10^5^ cells/well. After 12 hours, 1.5 µg all-in-one vector was transfected into each well of cells by lipo3000 (Thermo L3000015). After another 8 hours, 500 µL DMEM with 10% FBS was added to each well. Cells were collected 72 hours post-transfection.

Amplicon-seq was used to quantify the indel frequencies of each sgRNA. The gDNA was extracted from the collected cells using the Blood/Cells gDNA extraction kit from TIANGEN (TIANGEN DP304-03). The target sequences were enriched from the gDNA by two rounds of PCR amplification. The 1^st^ round of PCR was conducted in a 50 µL reaction, including 150 ng gDNA (∼1 µL), 250 nM forward primer, 250 nM reverse primer, 25 µL of Equinox Amplification Master Mix (2X) (WATCHMAKER GENOMICS 7K0014), and nuclease-free H2O. The PCR program was set as following: 45 s at 98 °C; 32 cycles of 15 s at 98 °C, 30 s at 60 °C, and 15 s at 72 °C; 2 mins at 72 °C; 4 °C hold. The 2nd round of PCR was conducted in a 50 µL reaction, including 2.5 µL products of the 1st round of PCR, 250 nM forward primer, 250 nM reverse primer, 25 µL of Equinox Amplification Master Mix (2X) (WATCHMAKER GENOMICS 7K0014), and nuclease-free H_2_O. The PCR program was set as following: 45 s at 98 °C; 12 cycles of 15 s at 98 °C, 30 s at 62 °C, and 15 s at 72 °C; 2 mins at 72 °C; 4 °C hold. AMPure XP beads (Beckman Coulter) were used to purify the PCR products, which were subjected to NGS sequencing. The primers used for the NGS library preparation was listed in **Supplementary Table 4**.

The Amplicon-Seq data was analyzed using CRISPResso 2.13 (--max_paired_end_reads_overlap 140 --min_paired_end_reads_overlap 10 --exclude_bp_from_left 0 --exclude_bp_from_right 0 --plot_window_size 40 --min_frequency_alleles_around_cut_to_plot 0.1)^24^.

### GUIDE-seq

To conduct GUIDE-seq, HEK293T cells were seeded into 12-well plate. At 90% confluency, cells were transduced by 25 pmol/well GUIDE-seq dsODN using Lipofectamine 3000. After 1 hour, cells were transduced by all-in-one vector, which expresses both SuperFi-Cas9 (or SpCas9) and sgRNA. (SuperFi-Cas9-tRNA-gRNA-all-in-one and spCas9-tRNA-gRNA-all-in-one). 72 hours post-transfection, cells were collected for gDNA extraction (TIANGEN DP304-03).

The extracted gDNA were fragmentized by Tn5 at 55 °C for 10 minutes. Each Tn5 fragmentation reaction includes 50 ng gDNA, 10 µL Tn5, 10 µL 5X Tn5 Digestion Buffer (50% DMF, 25 mM MgCl, 20 mM TAPS-NaOH (pH 8.5)), and H2O up to 50 µL.

The dsODN, Tn5 adaptors (Tn5 primeA and pegTn5+7N), and NGS primers (Tn5 PCR adaptor, Iguide forward, and Iguide reverse) used in GUIDE-seq were listed in **Supplementary Table 4**. GUIDE-seq sgRNA and target sequences were listed in **Supplementary Table 1**.

### AIdit-SuperFi architecture

AIdit-SuperFi is our own proposed deep learning model based on CNN and Transformer Encoder, which was specifically designed to predict the indel frequency of SuperFi-Cas9. As shown in Fig 4A, AIdit-SuperFi adopts a two-tower architecture, where the matching module and the context module shared the same parameter configs. In a bottom-to-up order, upon one-hot representation of input nucleotide sequences, each base module consists of CNN feature extraction layer with different kernel sizes, transformer encoder layer and mean pooling layer. The transformer encoder layer has 12 blocks, 12 attention heads in each block and an embedding dimension of 768. The left matching module is designed for modeling sgRNA spacer -NTS target pair and PAM (NGG), with pair lengths ranging from 19nt to 24nt. We removed PAM2-3 (GG) in on-target model as they remained unchanged across all sequences. The context module is designed for modeling upstream, target, PAM, and downstream sequences in NTS, with a total length of 63nt. The final prediction is regressed by a two-layer MLP module, using the sum of embeddings from the two base modules as input.

We consider a nucleotide pair that matches correctly between sgRNA and NTS as equivalent to the corresponding single nucleotide. During tokenization, the on-target matching vocabulary and the context vocabulary consist of 6 tokens: AA/A, CC/C, GG/G, UT/T, [MASK], and [PAD]. In contrast, the off-target matching vocabulary contains 26 tokens: AA/A, CC/C, GG/G, UT/T, AC, AG, AT, A-, -A, CA, CG, CT, C-, -C, GA, GC, GT, UA, UC, UG, U-, -T, [MASK], and [PAD]. Here, the symbol “-” represents insertion or deletion, the special token “[MASK]” is used for masking, and the special token “[PAD]” is used for padding.

Notably, instead of common word embedding, we introduced one-hot encoding and a CNN layer with a series of convolution kernels of different sizes for multiple granularity information extraction. All input sequences were padded to a predefined length, with the “[PAD]” token embedded as a zero vector. The output embeddings of all CNN kernels were concated and normalized before being fed into the following Transformer encoder module.

We replaced the absolute position encoding with Kernelized Relative Positional Embedding (KERPLE)^24^, where the kernel is defined as:

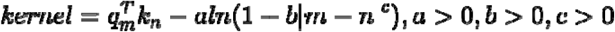

Here, a, b, and c are learnable parameters in each attention head of each layer. Additionally, we replaced the ReLU activation function with GEGLU^25^.

### Model pretraining and fine-tuning

We used high throughput screening data of SuperFi-Cas9 profiling library L2 and L4 for training. L2, with 322,530 samples, was used for on-target models. L4, with 100,257 samples, including matched sgRNA with different PAM, single mismatch, double mismatch, single insertion, and single deletion, was used for off-target models. In addition to total indel frequency, on-target dataset also records frameshift indel frequency and non-frameshift indel frequency, which were used to train our frameshift and non-frameshift models. We randomly split each dataset into training, validation, and test sets with a ratio of 8:1:1. To avoid data leakage, samples from the same gene target will be divided into the same set **(Supplementary Table 5)**.

To enhance the performance of the model, each base module was pretrained separately using a Masked Language Modeling (MLM) objective (sFig. 10C) with 15 percent of tokens randomly masked in each sequence. Of the masked tokens, 80% were replaced by special token “[MASK]”, 10% were replaced by random token and 10% were left unchanged.

Initially, the models were pretrained on the human genome assembly GRCh38.p14 (https://www.ncbi.nlm.nih.gov/datasets/genome/GCF_000001405.40/), with repeats and unannotated regions removed. 40,960,000 sequences, with lengths ranging from 64 to 512, were randomly selected. After that, the models’ pretraining was continued using the corresponding downstream training set. Duplicate sequences in the context training set were also removed.

The models were then fine-tuned to predict the total on-target indel frequency (“on-target”), the on-target frameshift indel frequency (“frameshift”), the on-target non-frameshift indel frequency (“non-frameshift”) and the off-target indel frequency ("off-target"). The Mean Squared Error (MSE) is chosen as the loss function. Both the pretraining and fine-tuning processes utilized the AdamW^26^ optimizer with β_1_ = 0.9, β_2_ = 0.999, D = 1e-8, and a weight decay of 0.01. Other hyperparameters for the pretraining and fine-tuning tasks are provided in the **(Supplementary Table 6)**.

### Baseline models

We trained multiple deep learning baseline models, all of which adopted the same two-tower architecture, replacing the CNN layer and transformer encoder layers with other encoders (sFig. 10B). These encoders included a convolutional neural network (CNN) (sFig. 10E), a recurrent neural network (RNN) (sFig. 10F), a multilayer perceptron (MLP) (sFig. 10G), and the pretrained models Caduceus-PS^27^ and DNABERT-2^28^. L2 and L4 were also used to train these models. Given that these pretrained models could only accept single-stranded nucleic acid as input, we were unable to train off-target models with them. Except for DNABERT-2, which had a learning rate of 1e-5, all other hyperparameters were identical to those in AIdit-SuperFi fine-tuning.

We also established baselines using conventional machine learning algorithms, including XGBoost (XGB)^29^, Random Forest (RF)^30^, and ElasticNet^31^ (sFig. 10D). Each nucleotide pair type or single nucleotide type at every position of matching and context was treated as a feature. The training set and validation set were merged, and 5-fold cross-validation along with random search were employed to select the optimized parameter combination.

For XGBoost, the n_estimators parameter was selected from the range of 100 to 5000, the max_depth parameter from 3 to 20, the learning_rate parameter from 0.01 to 0.1, the subsample parameter from 0.5 to 1, the colsample_bytree parameter from 0.5 to 1, the gamma parameter from 0 to 0.5, the reg_alpha parameter from 0 to 1, and the reg_lambda parameter from 0 to 1.

For Random Forest, the n_estimators parameter was chosen from 50 to 2000, the max_depth parameter from 3 to 20, the min_samples_split parameter from 2 to 100, the min_samples_leaf parameter from 1 to 50, and the max_features parameter from 0.5 to 1. For ElasticNet, the alpha parameter was selected from 0.01 to 100, and the l1_ratio parameter from 0 to 1.

### SuperFi-Cas9 protein expression and purification

SuperFi-Cas9 was expressed in Escherichia coli BL21(DE3) cells. The cells were grown at 37°C in LB medium until the OD600 reached 0.6-0.8, followed by protein expression induced with 0.2 mM IPTG at 16°C for 13 hours. Cells were harvested by centrifugation and resuspended in lysis buffer containing 500 mM NaCl, 20 mM HEPES pH 7.4, and 10% glycerol. The cells were lysed using a microfluidizer (ATS Scientific Inc) operated at 600 bars. The lysate was then subjected to sonication (40% amplitude, 10 seconds pulse on, 30 seconds pulse off for 20 cycles) for 30 minutes to ensure efficient DNA removal. The lysate was clarified by centrifugation at 17,000g for 30 minutes. The protein was purified using TALON cobalt affinity resin. The clarified lysate was applied to the pre-equilibrated TALON resin with lysis buffer, washed extensively with 20 mM imidazole-supplemented lysis buffer, and the protein was eluted with 300 mM imidazole-supplemented lysis buffer. The eluted protein showed minimal DNA contamination with an A260/280 ratio of 0.79. The protein was further purified by size exclusion chromatography using a Superdex S200 Increase 10/300 column equilibrated with a buffer containing 150 mM NaCl, 20 mM HEPES pH 7.4.

### Nucleic acid preparation

Three synthetic sgRNAs of different spacer lengths were chemically synthesized (GenScript) and dissolved in nuclease-free water to a stock concentration of 10 mM. The target DNA duplex was prepared using two complementary oligonucleotides. The oligonucleotides were dissolved in nuclease-free water to 10 mM stock solutions, mixed in equimolar ratios, denatured at 95°C, and annealed by gradual cooling to room temperature to form the target dsDNA. The synthetic sgRNAs carry 2’-O-methyl and phosphorothioate modifications at the first three 5’ and 3’ terminal RNA residues (**Supplementary Table 7**).

### Ternary complex formation of SuperFi-Cas9 with sgRNA and dsDNA

To form the ternary complex, complementary DNA oligonucleotides were first denatured at 95°C and annealed by gradual cooling to room temperature. The SuperFi-Cas9 was incubated with sgRNA at a molar ratio of 1:1.5 for 10 minutes at room temperature, followed by the addition of annealed target dsDNA (molar ratio of 1:2.3 relative to Cas9) for an additional 10 minutes. The final molar ratio of SuperFi-Cas9:sgRNA:dsDNA in the complex was 1:1.5:2.3. Complex formation was verified by size exclusion chromatography using a Superdex S200 Increase 10/300 column equilibrated with a buffer containing 150 mM NaCl, 20 mM HEPES pH 7.4, 10 mM MgCl2, and 0.5 mM TCEP. Successful compl×ex assembly was confirmed by both a shift in elution volume and changes in the A260/280 ratio compared to the individual components (sFig. 20). The SEC fractions containing ternary complex of SuperFi-Cas9 were pooled for sample concentration. Given that the SuperFi-Cas9 with sgRNA-dsDNA complex was pre-formed, protein quantification was not determined accurately. Instead, combined fractions were concentrated approximately 10-fold from their initial absorbance of 60 mAU. Similar concentration ratios were applied across all sample preparations to ensure consistent relative concentrations.

### Cryo-EM grid preparation and data collection

For cryo-EM grid preparation, 3 μL of the concentrated sample was applied to glow-discharged (PELCO easiGlow, 15mA for 50s) Quantifoil Cu 400 mesh R1.2/1.3 grids. Vitrification was performed using a Vitrobot Mark IV (Thermo Fisher Scientific) with the following parameters: temperature at 4°C, 100% humidity, blot force 1, single blot, 5 seconds wait time, 4 seconds blotting time, and 0 seconds drain time. The grids were plunge-frozen in liquid ethane cooled by liquid nitrogen. Data collection was carried out on a Titan Krios microscope (Thermo Fisher Scientific). The detailed information and parameters of data collection are listed in **Supplementary Table 8**.

### Cryo-EM data processing

The final reconstruction parameters and particle statistics are summarized in **Supplementary Table 8**. The collected cryo-EM movie stacks were processed for beam-induced motion correction and dose-weighting using MotionCor2^32^. Micrograph defocus estimation was performed using CTFFIND4^33^. Initial image processing was conducted using the SAMUEL (Simplified Application Managing Utilities of EM Labs) package^34^. The dose-weighted micrographs underwent 3x binning using "sampilcopy2d.py", followed by manual screening. Automated particle selection was performed using "samautopick.py", followed by stack generation with "sampilboxparticle.py". Initial 2D classification was carried out using "samtree2dv3.py". The particle data sets were further refined through additional rounds of 2D and 3D classification in RELION 2.0^35^ and RELION 3.0^36^ to exclude bad particles. Multiple processing strategies were employed for map refinement using CryoSPARC^37^, as detailed in **Supplementary Table 8.**

### 3D classification of SuperFi-Cas9:20nt sgRNA complex in cryoSPARC

To identify both conformational heterogeneity and potential redundant particle populations in the SuperFi-Cas9 with 20nt sgRNA dataset, additional classification was performed using the "3D Classification" tool in CryoSPARC^37^. A total of 242,591 particles were subjected to 3D classification with a K value of 6 and a mask encompassing the entire complex (**Supplementary Table 8)**. The classification parameters were set as follows: target resolution at 6 Å, EM learning rate initialization at 0.75, O-EM learning rate half-life at 100%, auto solvent mask near parameter at 20 Å, auto solvent mask far parameter at 30 Å, and class similarity at 0.25. After 96 iterations, five distinct classes emerged. Each class was independently refined using the "Non-Uniform Refinement" and "Local Refinement" tools in CryoSPARC. Four of these classes corresponded to the previously identified Classes A-D, while one class containing 52,037 particles exhibited a unique conformation and was designated as Class E.

### Structural model building

Predicted structures of AlphaFold 3^38^ were used as starting models. Model adjustment and preliminary refinement were completed using COOT^39^. Atomic model refinement was completed using Phenix^40^. All structural figures were generated using ChimeraX^41^.

## Acknowledgments

We thank the Torch Project team and Dr. Maofu Liao for their invaluable assistance with Cryo-EM sample preparation and data collection. We thank the Biomedical Research Core Facilities, Laboratory Animal Resource Center, and High-Performance Computer Center of Westlake University for their excellent technical assistance. We are thankful for the inspiring suggestions from Dr. Huaizong Shen, Dr. Dan Ma, Dr. Ruixue Wang, Dr. Haishan Gao from Westlake University and Dr. Yanli Wang from the Institute of Biophysics (Chinese Academy of Sciences) on analyzing and interpreting the Cryo-EM data. We also appreciate the feedback Dr. Dangsheng Li provided during the development of this study. This research was supported by the Pioneer and Leading Goose R&D Program of Zhejiang (grant no. 2024SSY0033).

## Author contributions

L. Ma and Z. Lu conceived the project and designed the experiments. Z. Lu designed the profiling library. R. Wei conducted the experiments with help from J. Li, X. Wang, Y. Wei, and H. Zhang. Z. Lu, Q. Zhang, and R. Zheng conducted the bioinformatics analysis. R. Zheng built the deep learning models with help from H. Chen, Z. Lu, and Q. Zhang. Y. Shin, B. Liu, and Y. Chen collected Cryo-EM data. R. Zheng and Y. Shin processed and build model for the Cryo-EM data. Q. Zhang and H. Chen built the GUI of AIdit-SuperFi prediction model. L.M. wrote the manuscript with input from all co-authors.

## Data availability

Data deposit:

NGS data is available from GEO GSE279806. Please use the following token to access: axslkqgqbzedxsh

CryoEM data is available from PDB (Supplementary Table 8).

---

**sFig. 1.**
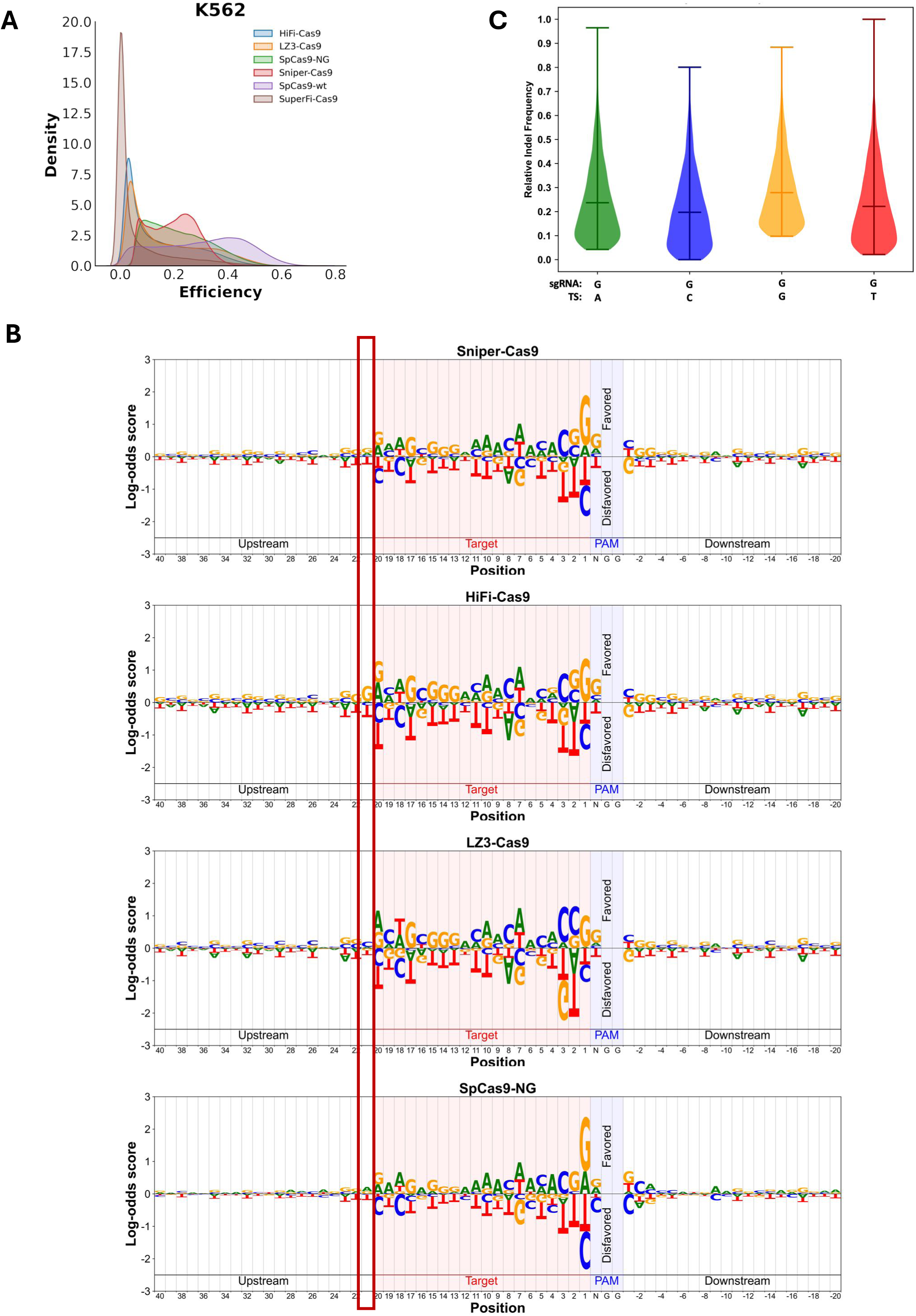

**sFig. 2.**
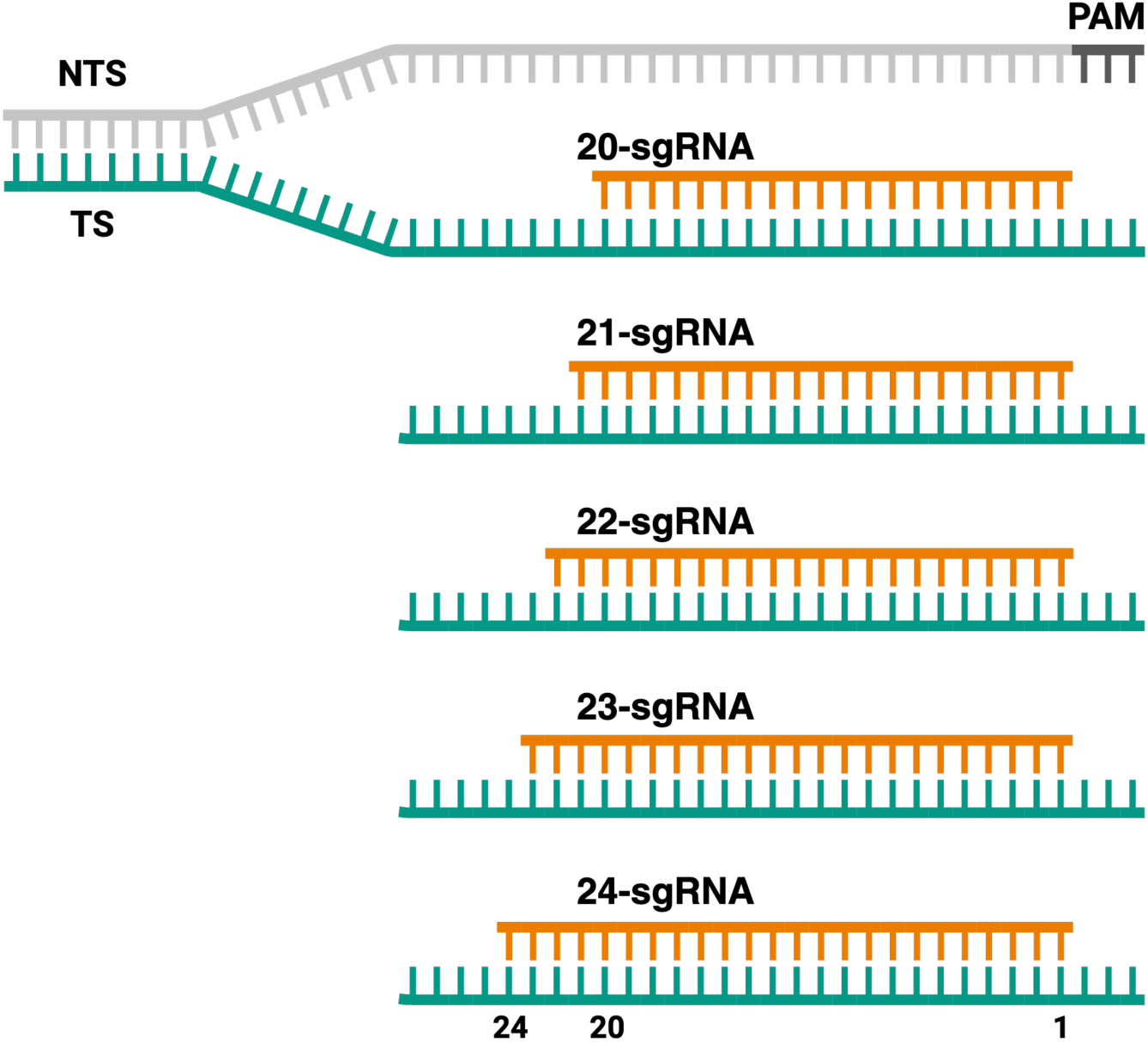

**sFig. 3.**
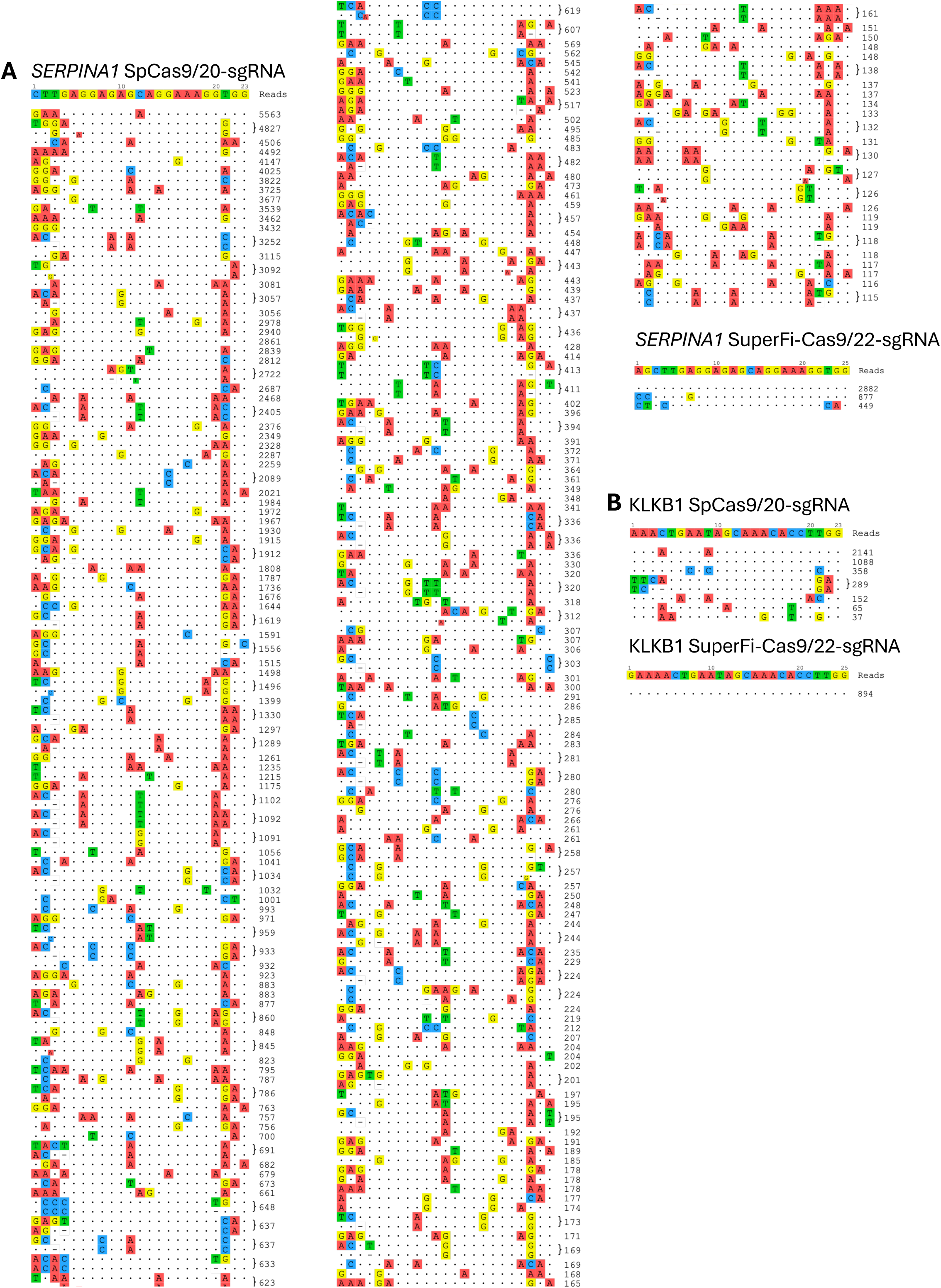
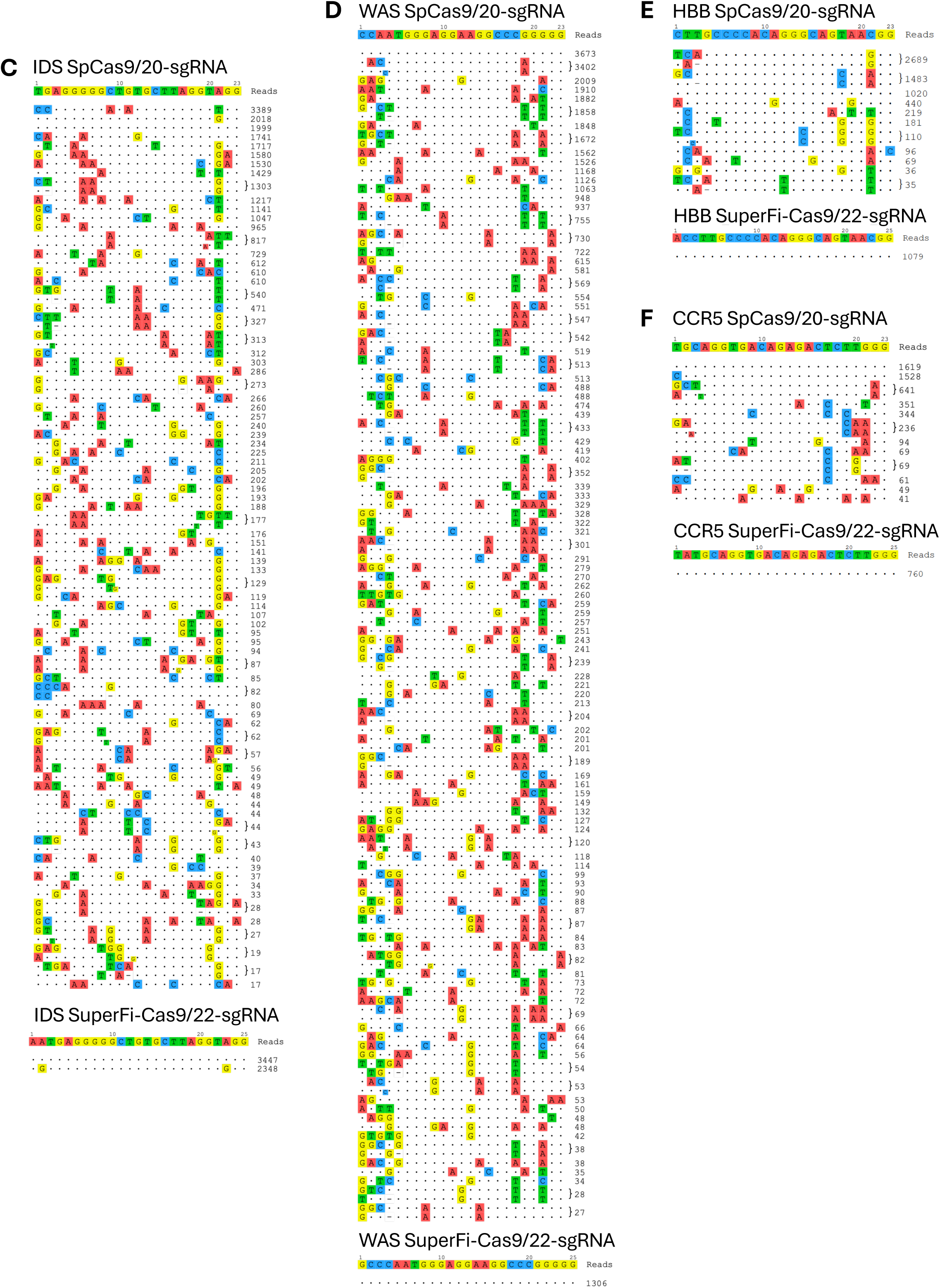

**sFig. 4.**
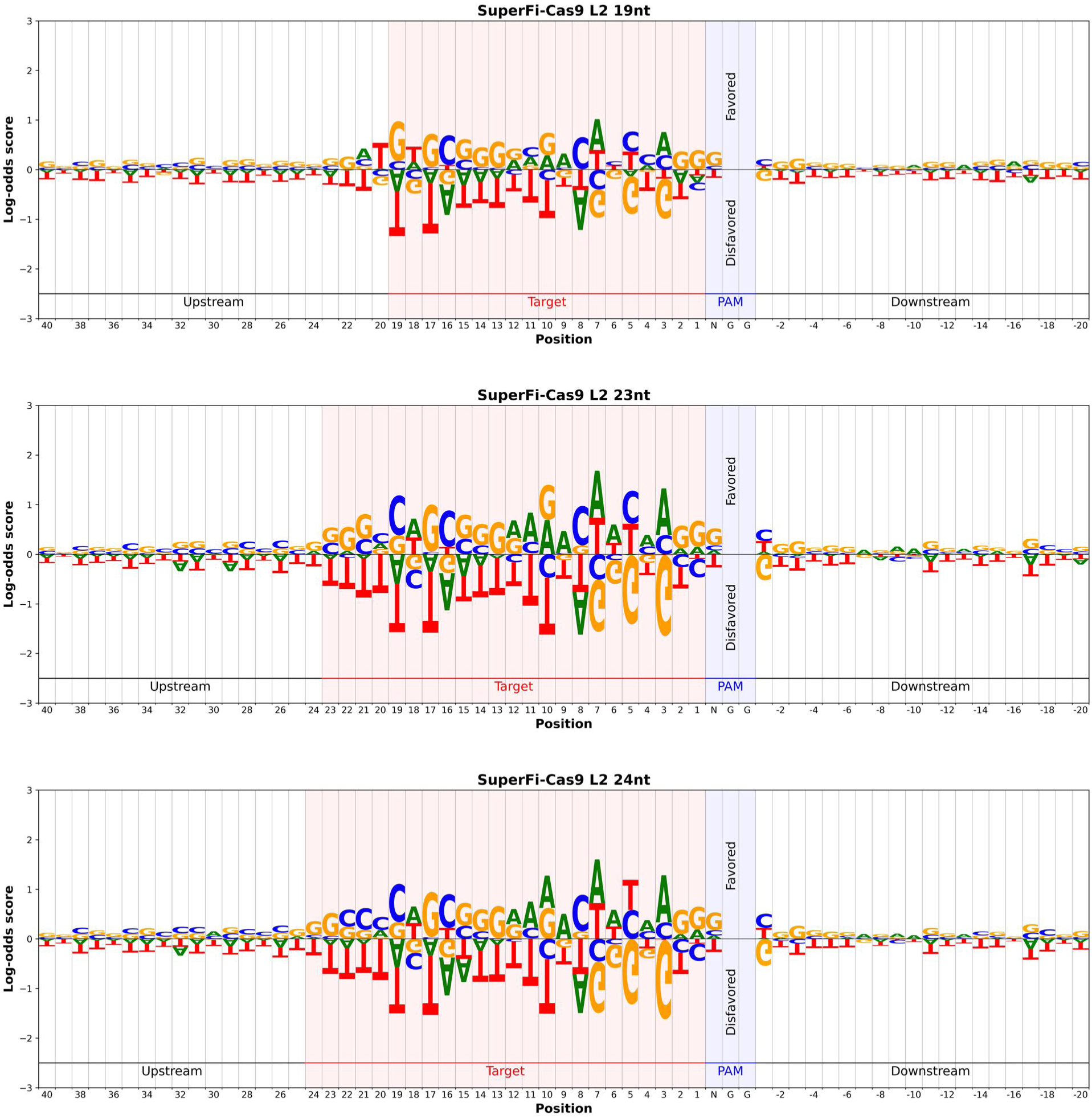

**sFig. 5.**
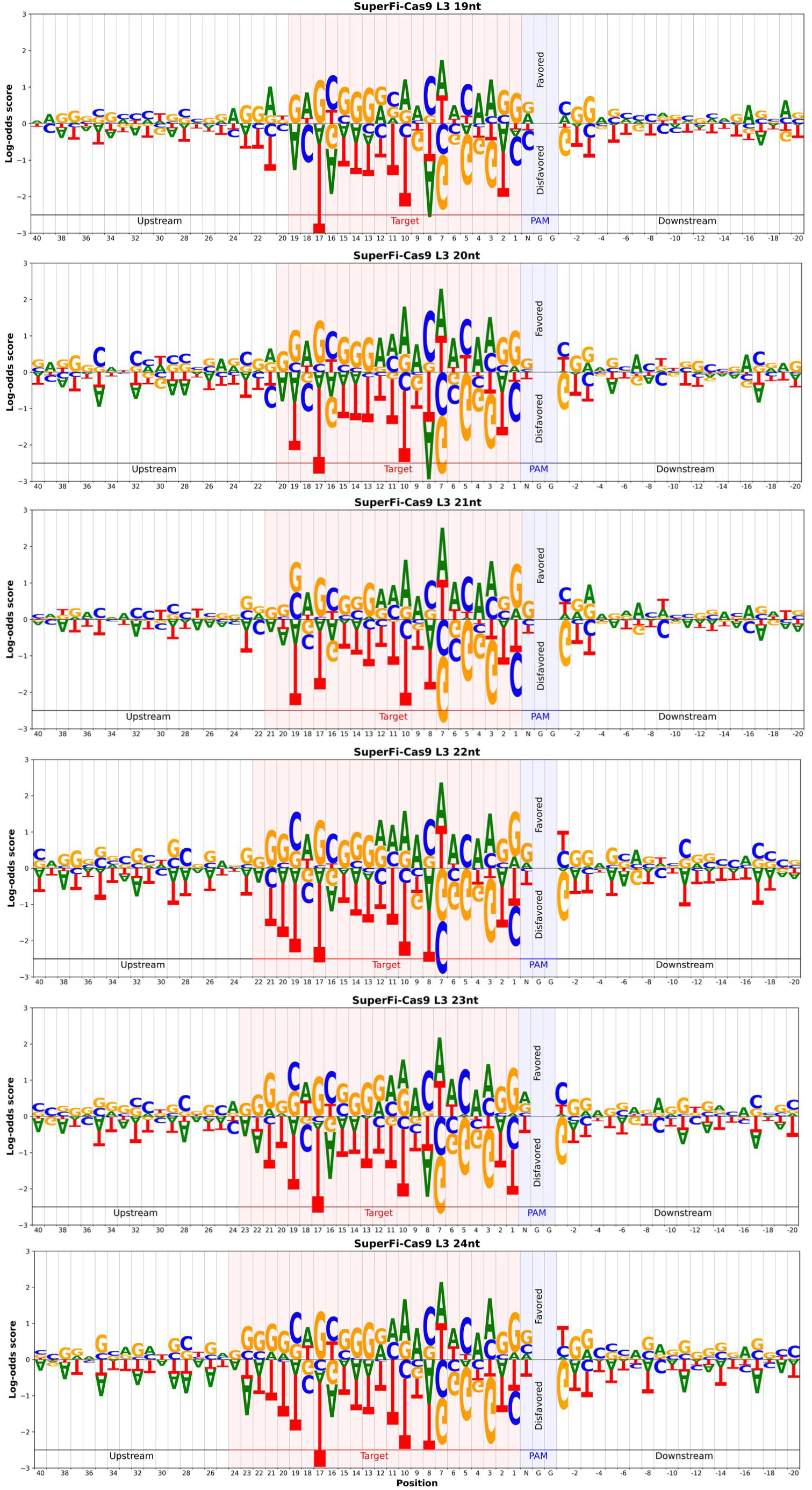

**sFig. 6.**
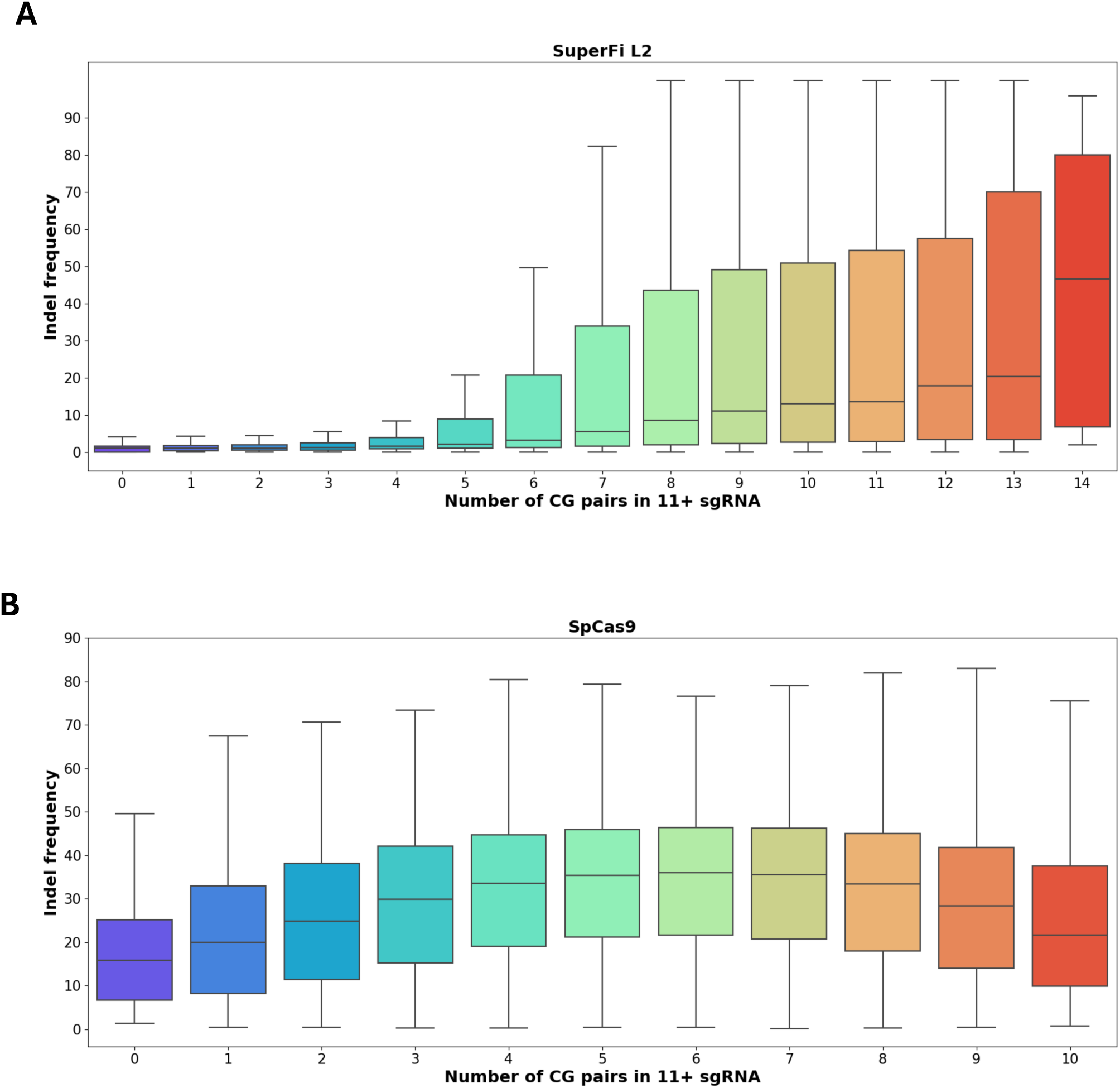

**sFig. 7.**
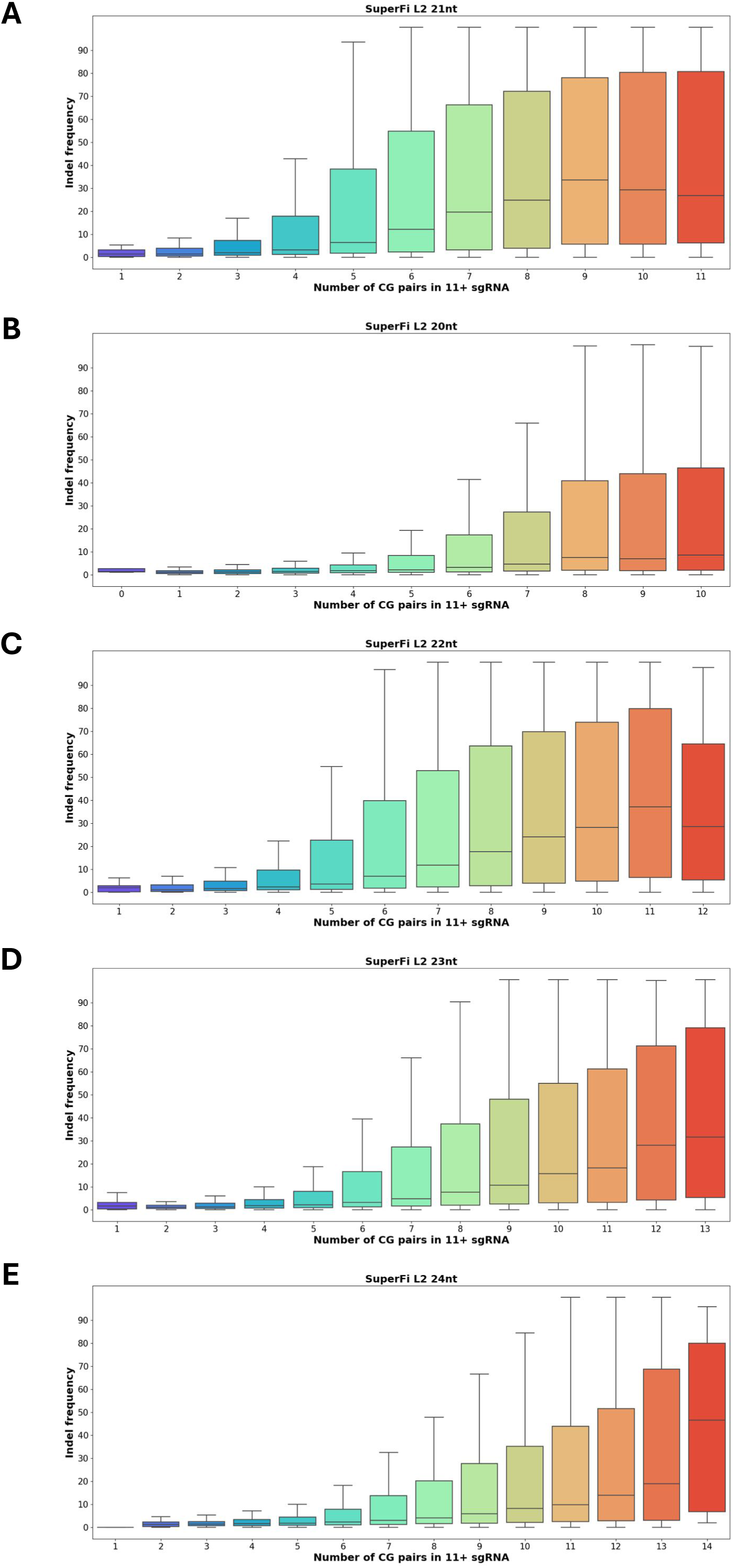

**sFig. 8.**
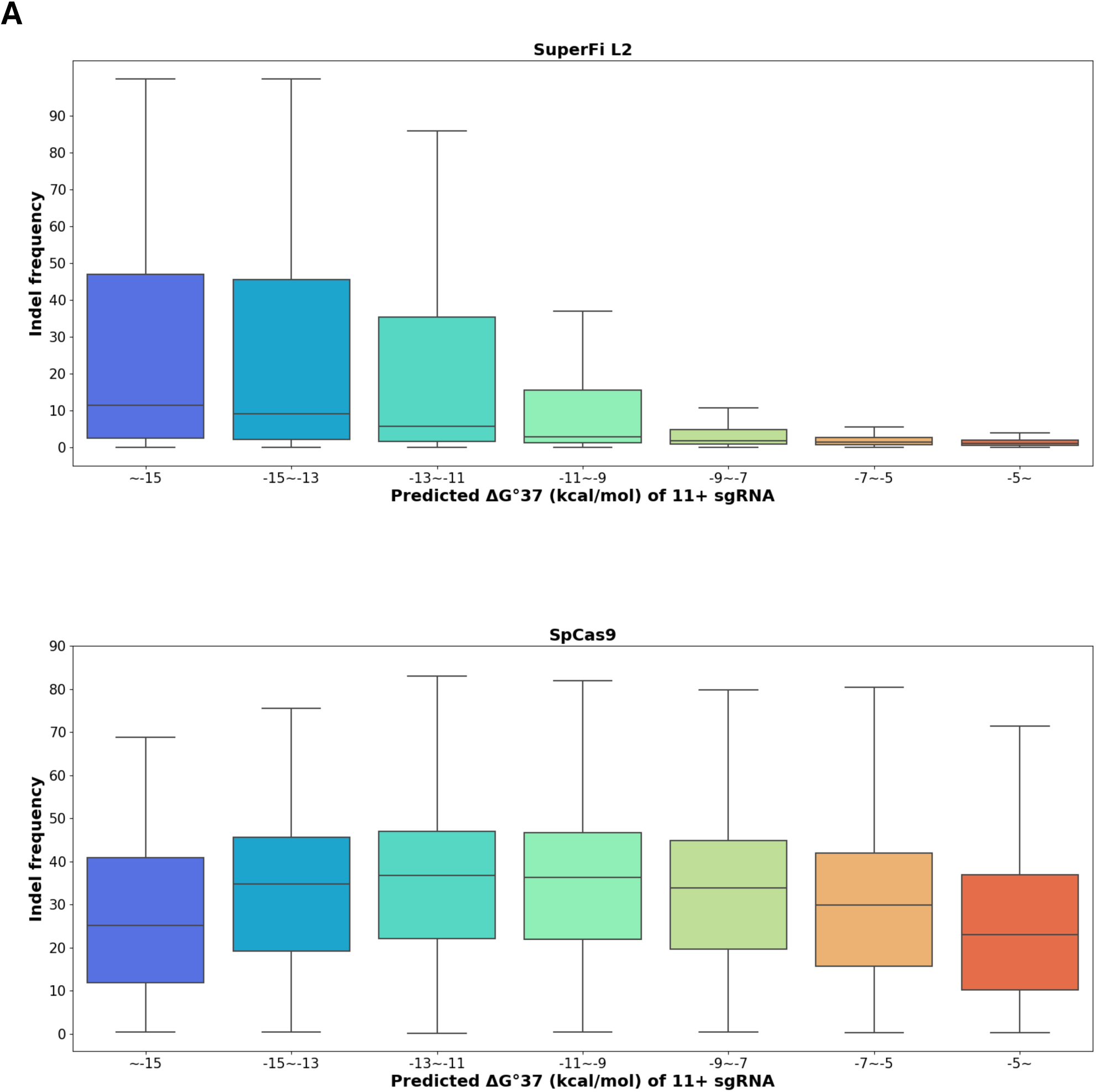

**sFig. 9.**
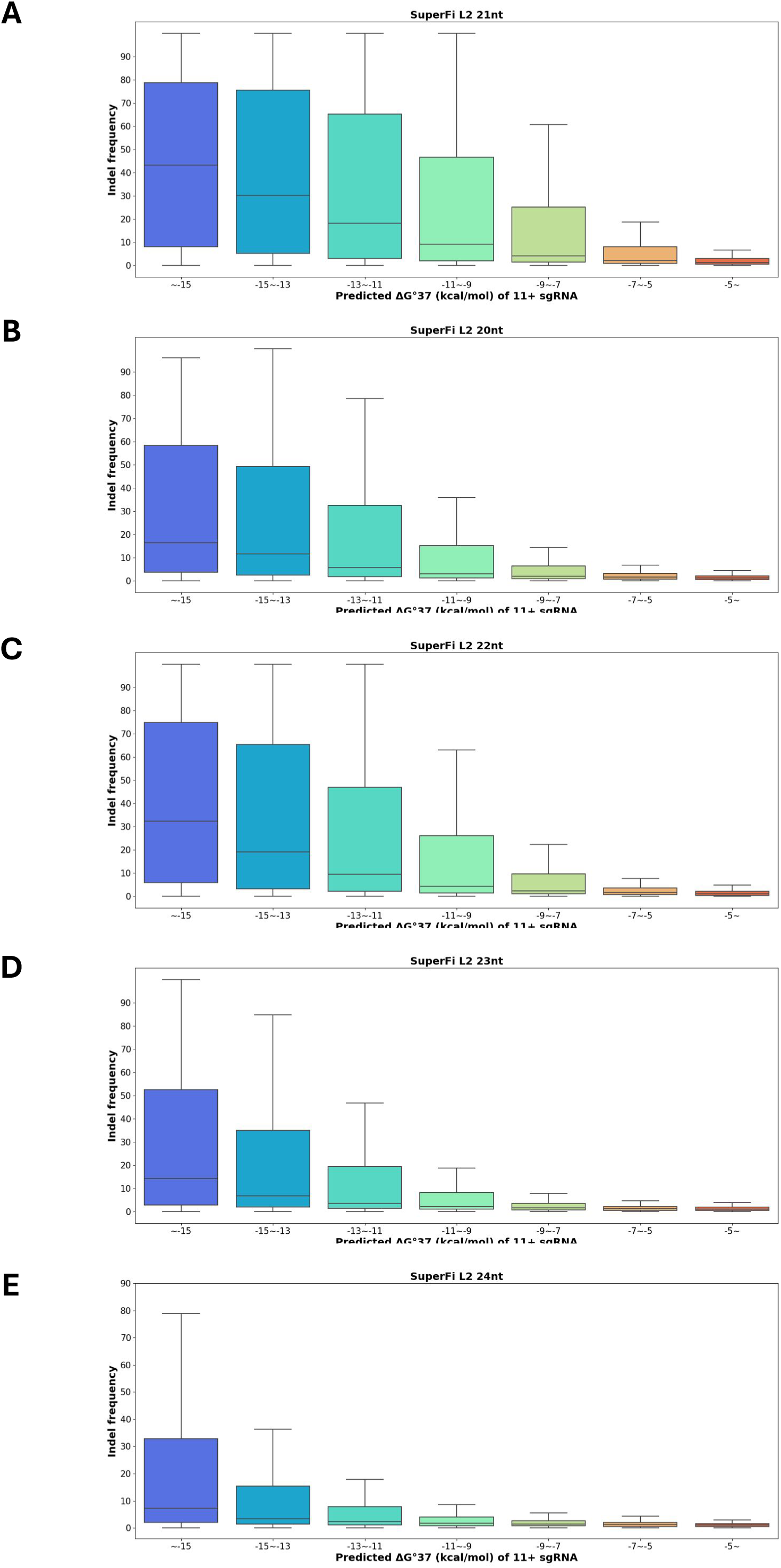

**sFig. 10.**
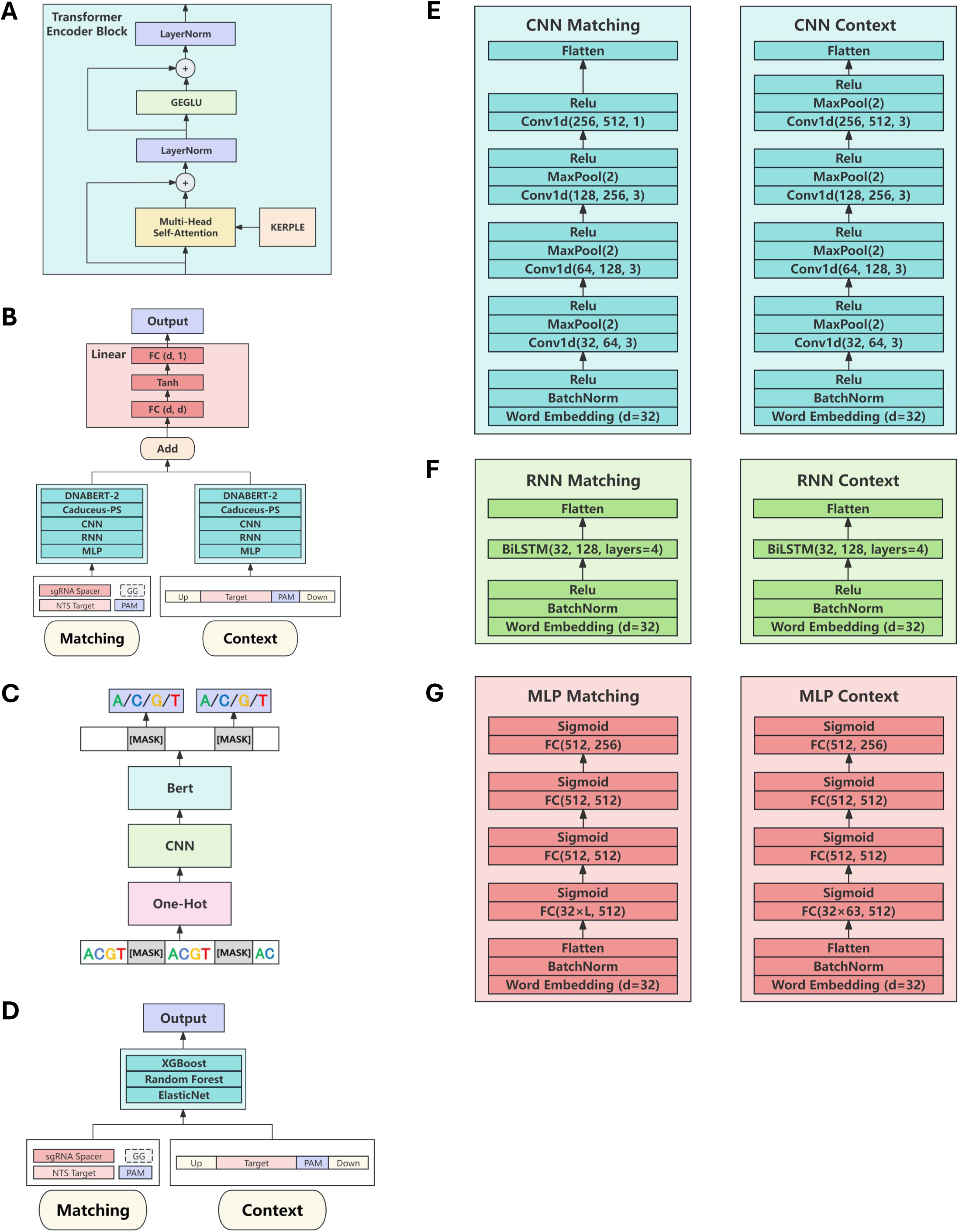

**sFig. 11.**
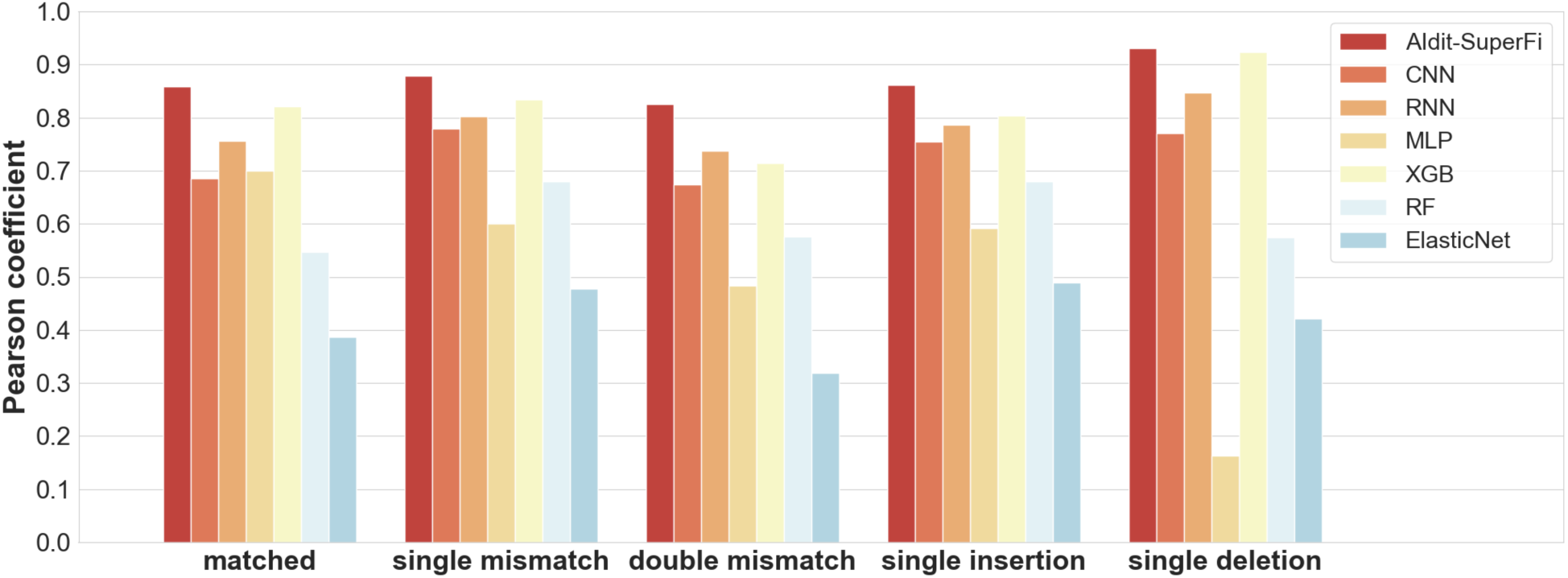

**sFig. 12.**
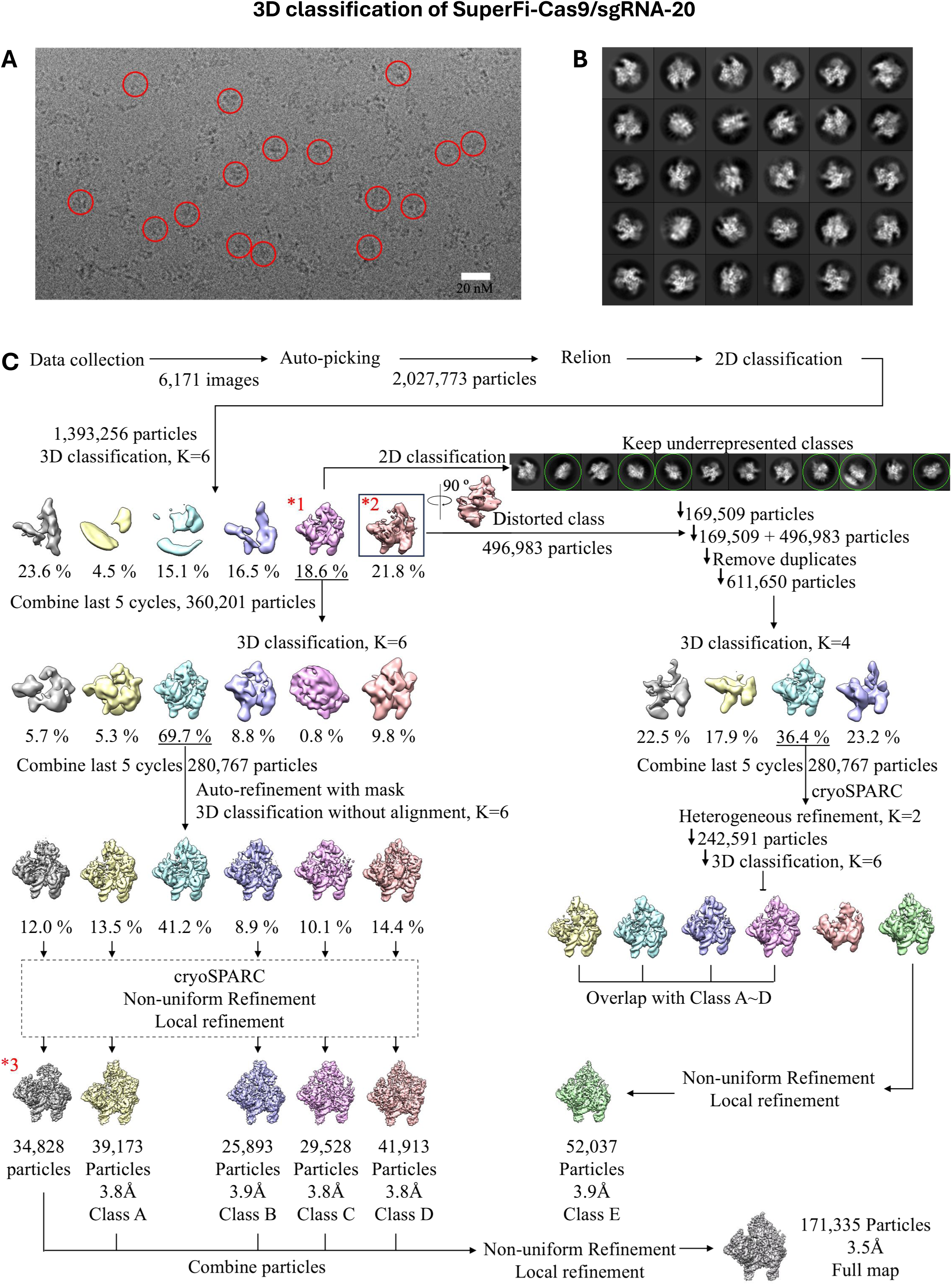
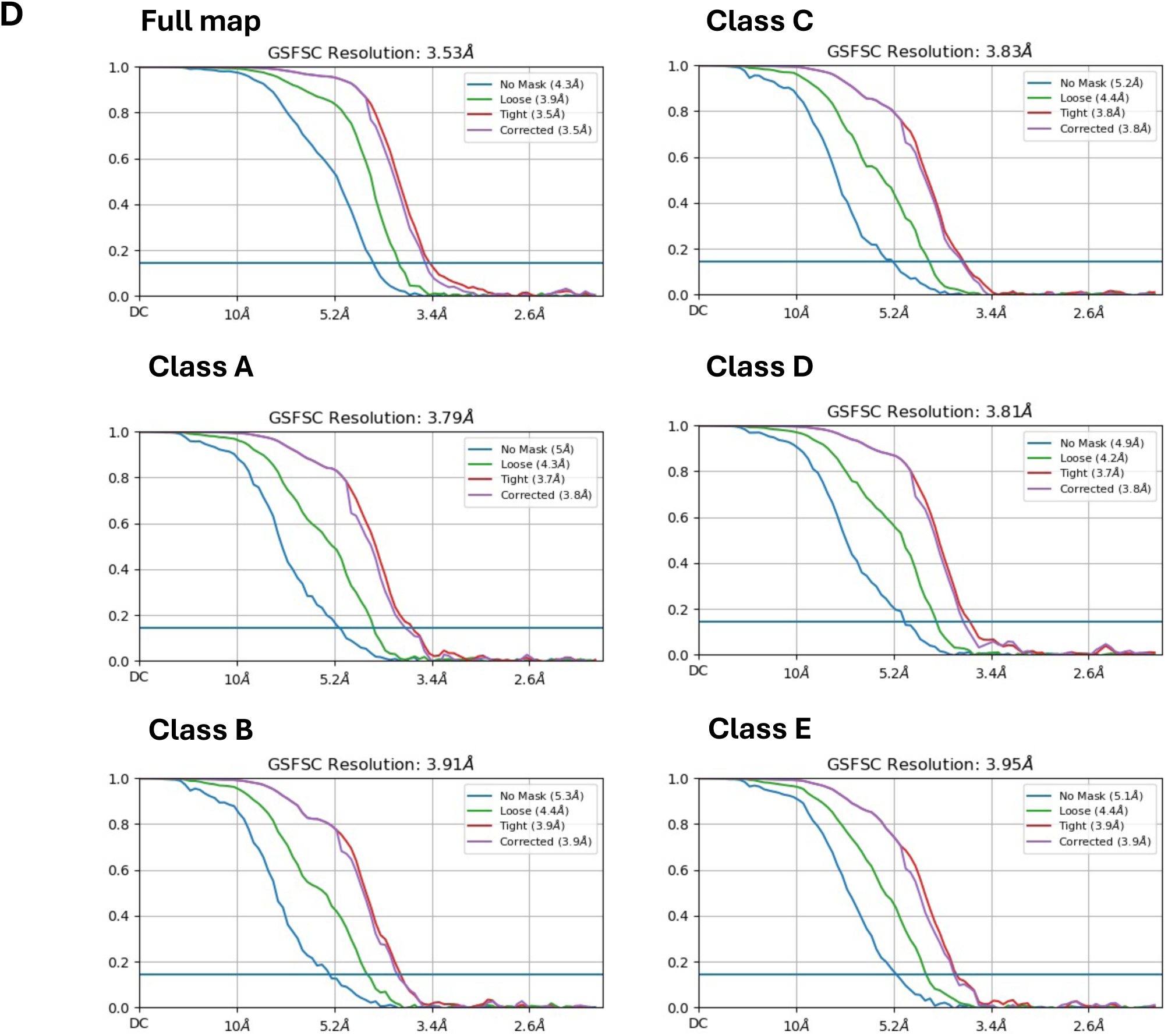
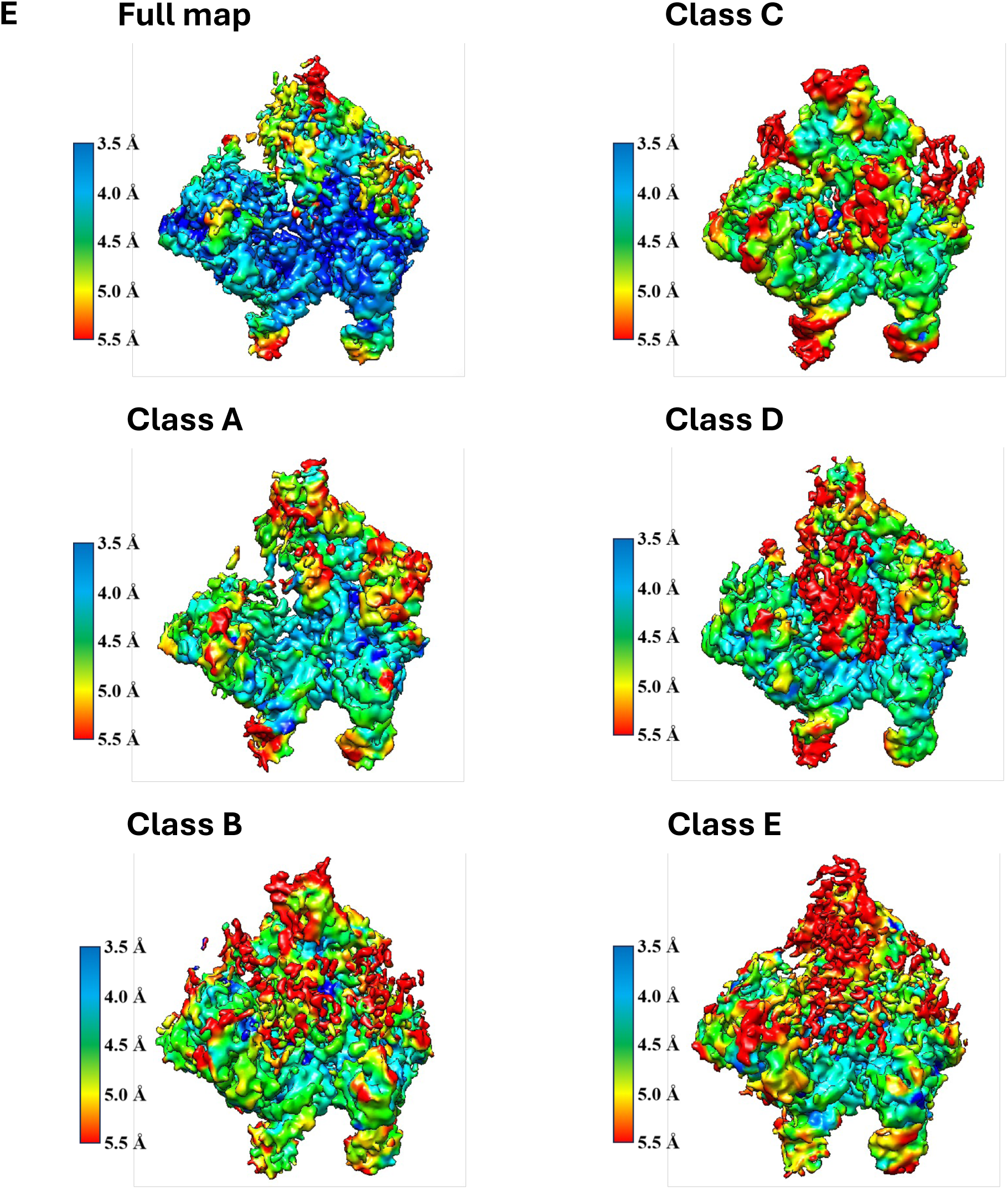

**sFig. 13.**
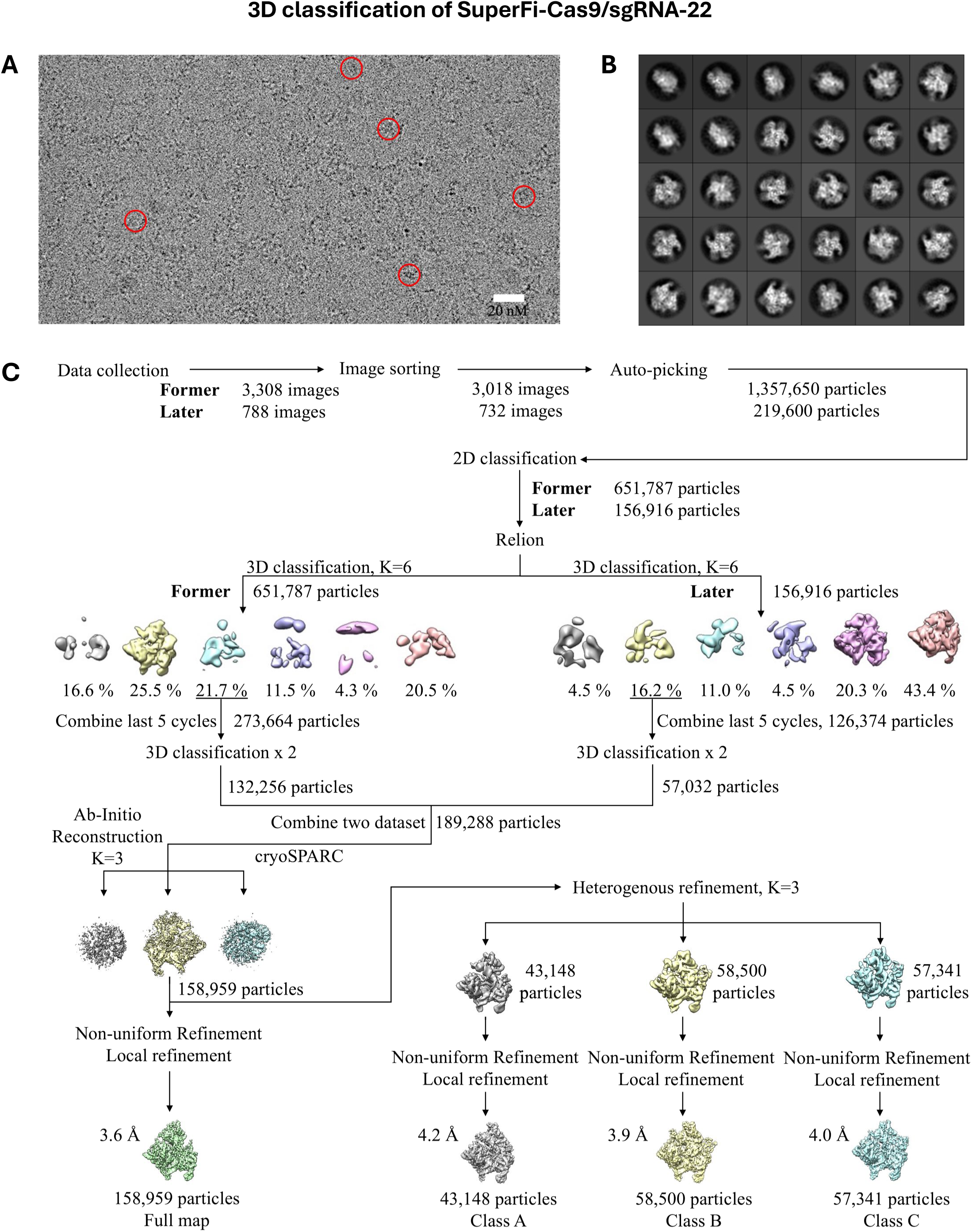
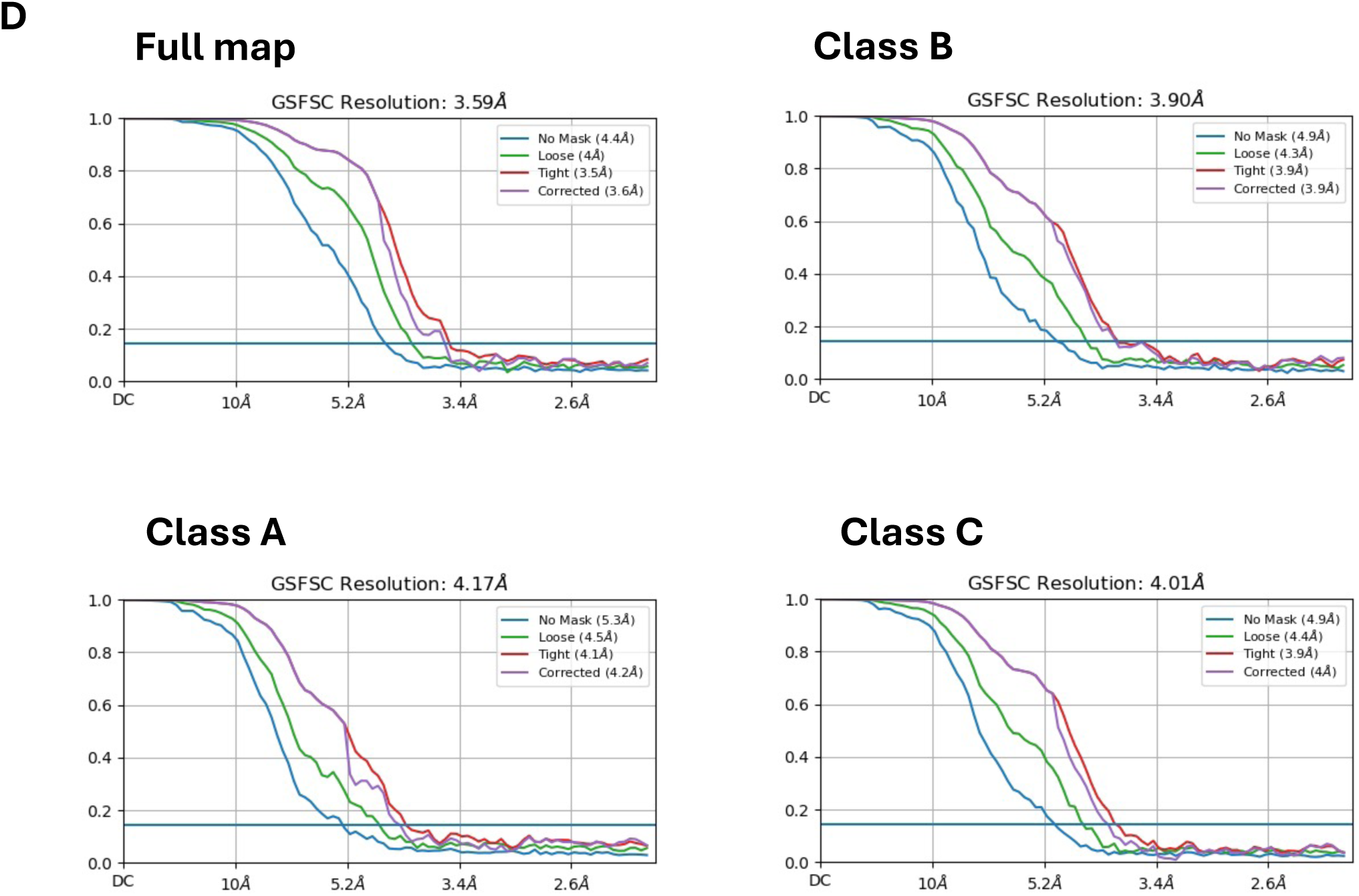
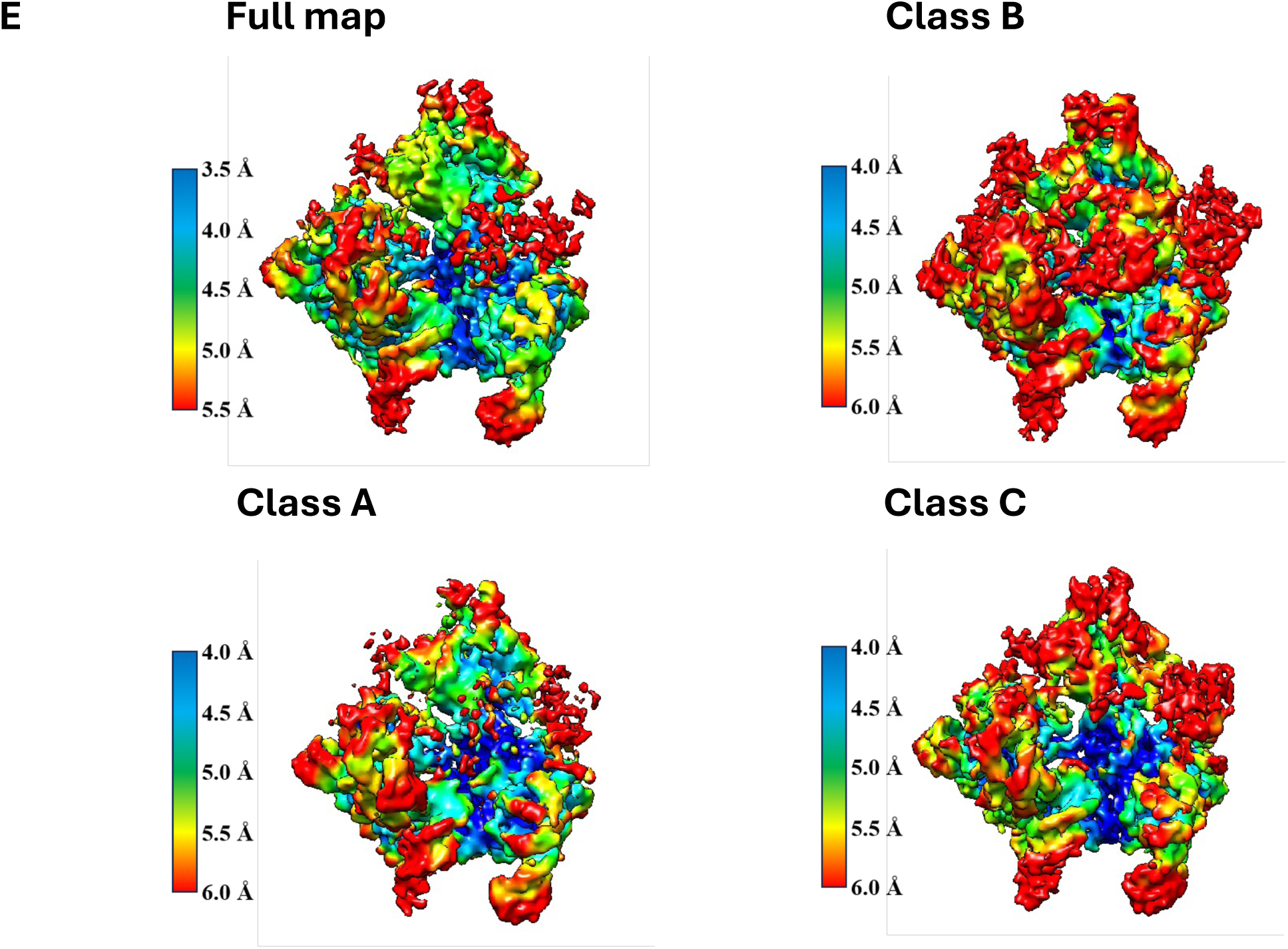

**sFig. 14.**
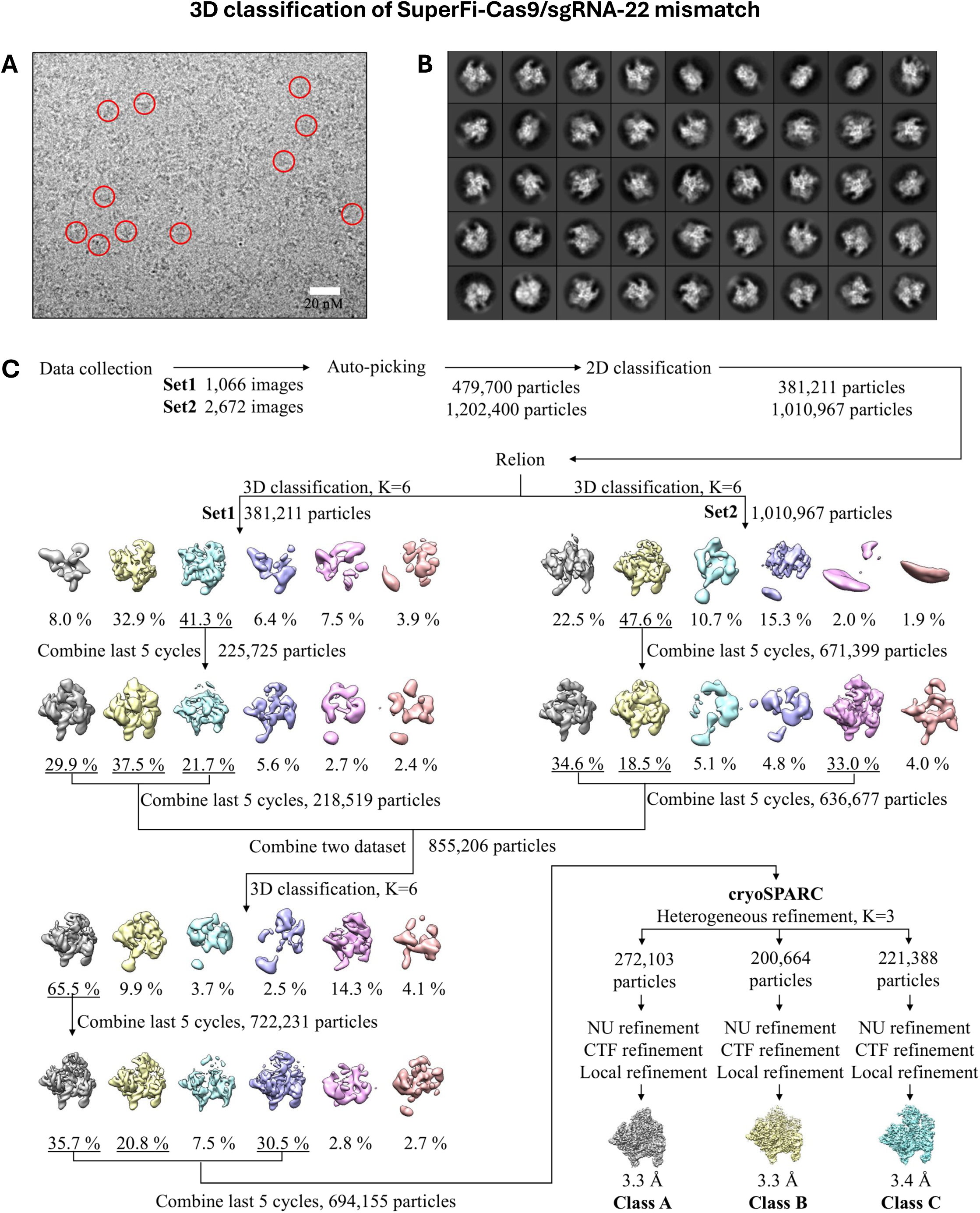
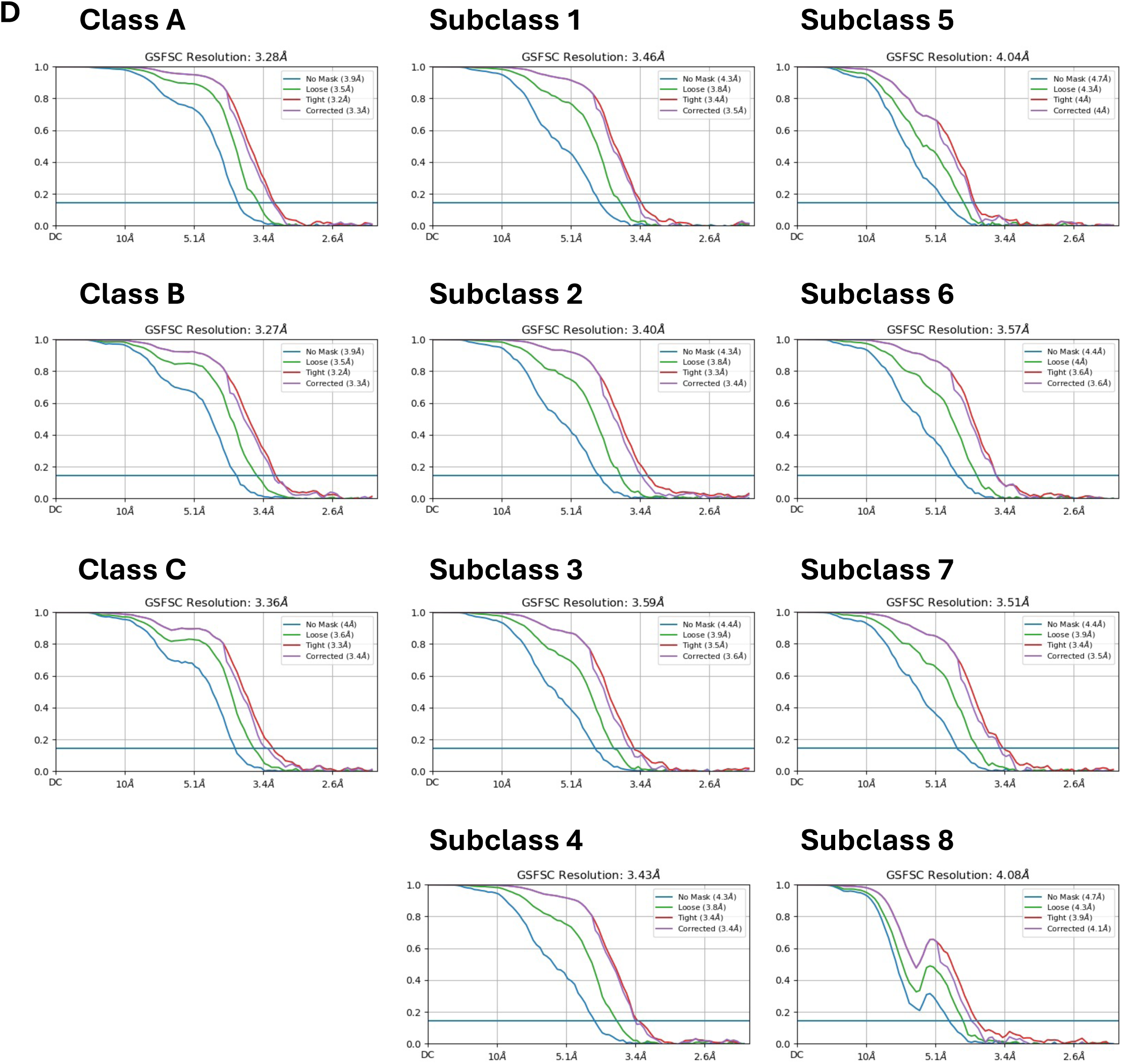
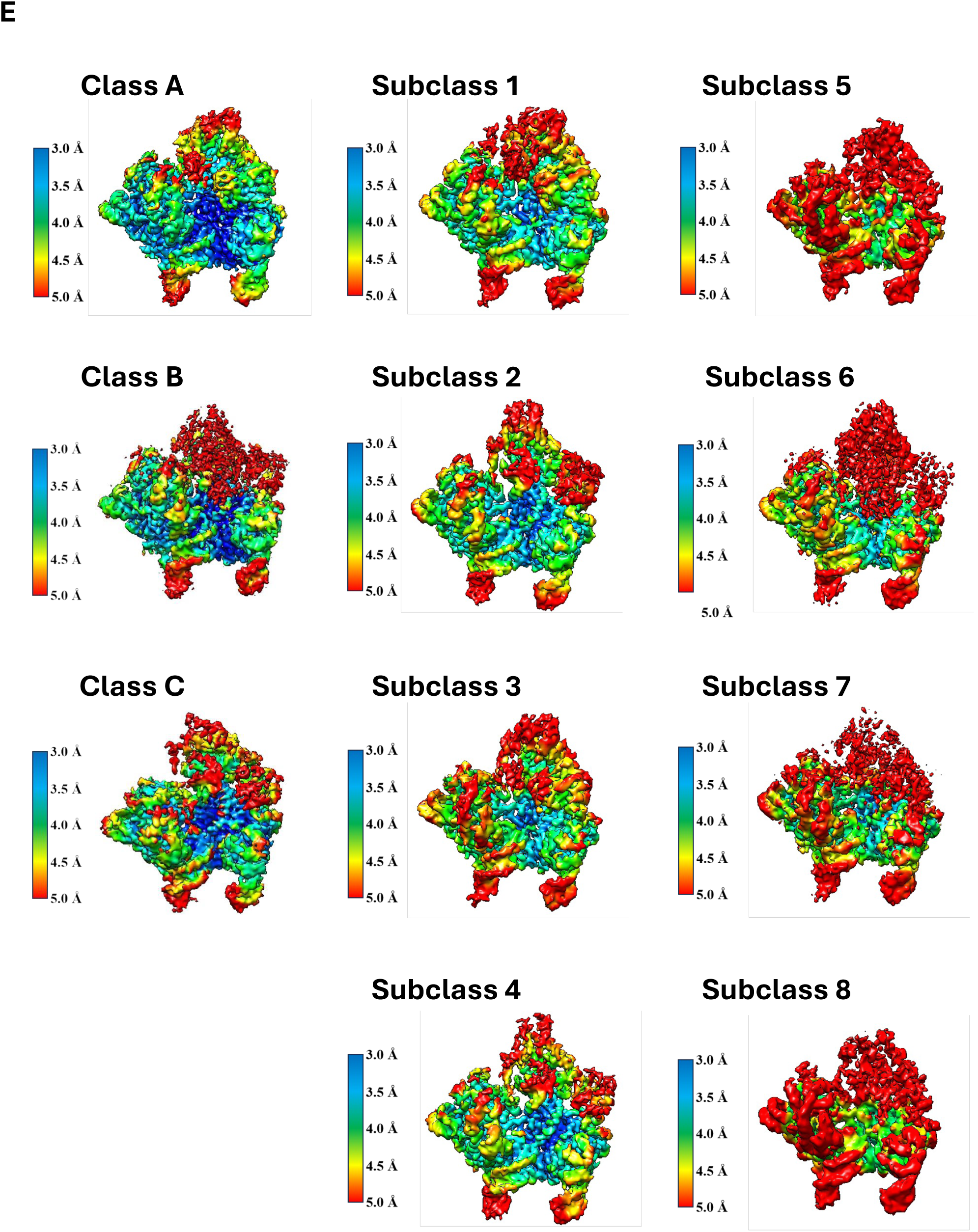

**sFig. 15.**
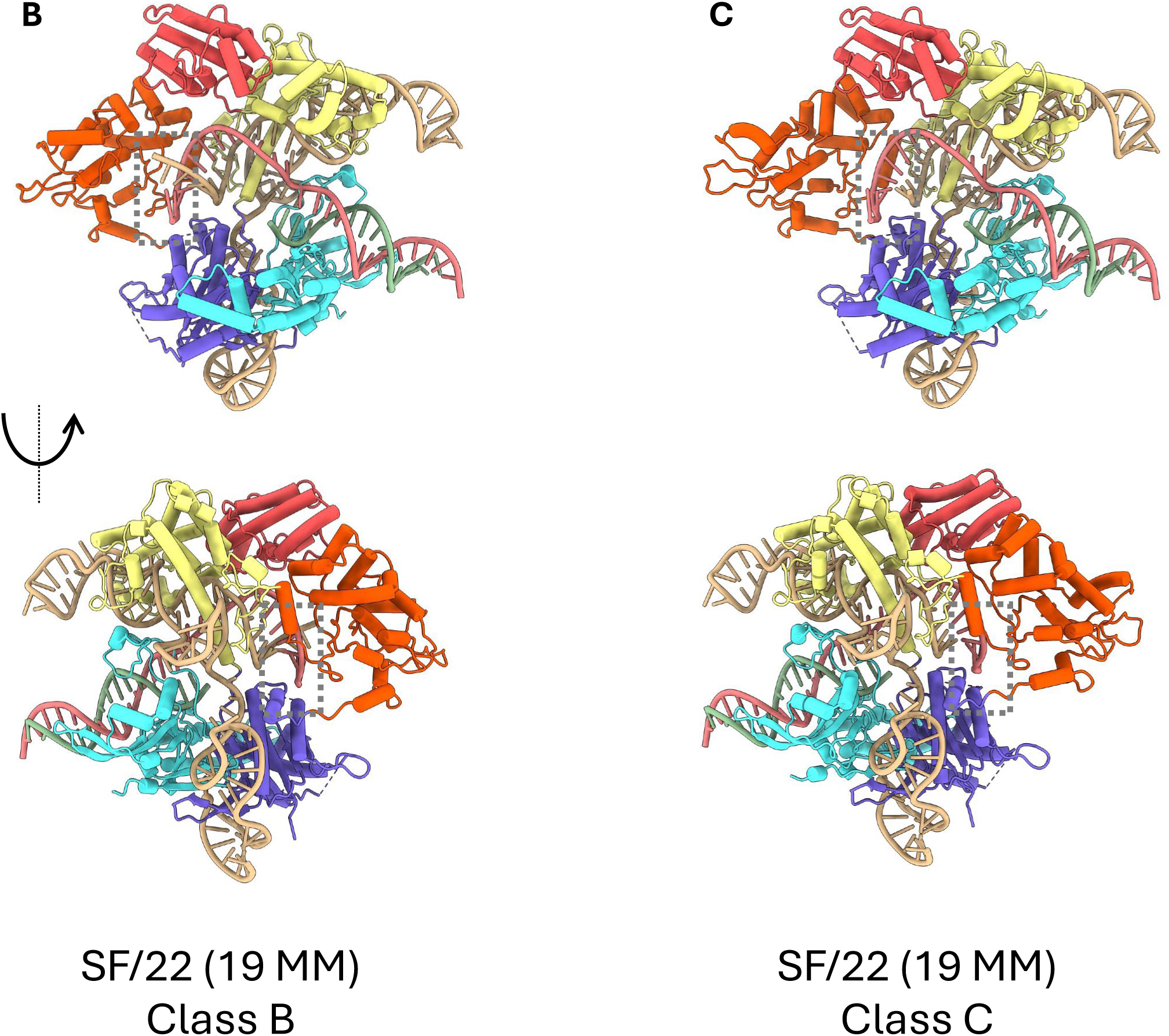

**sFig. 16.**
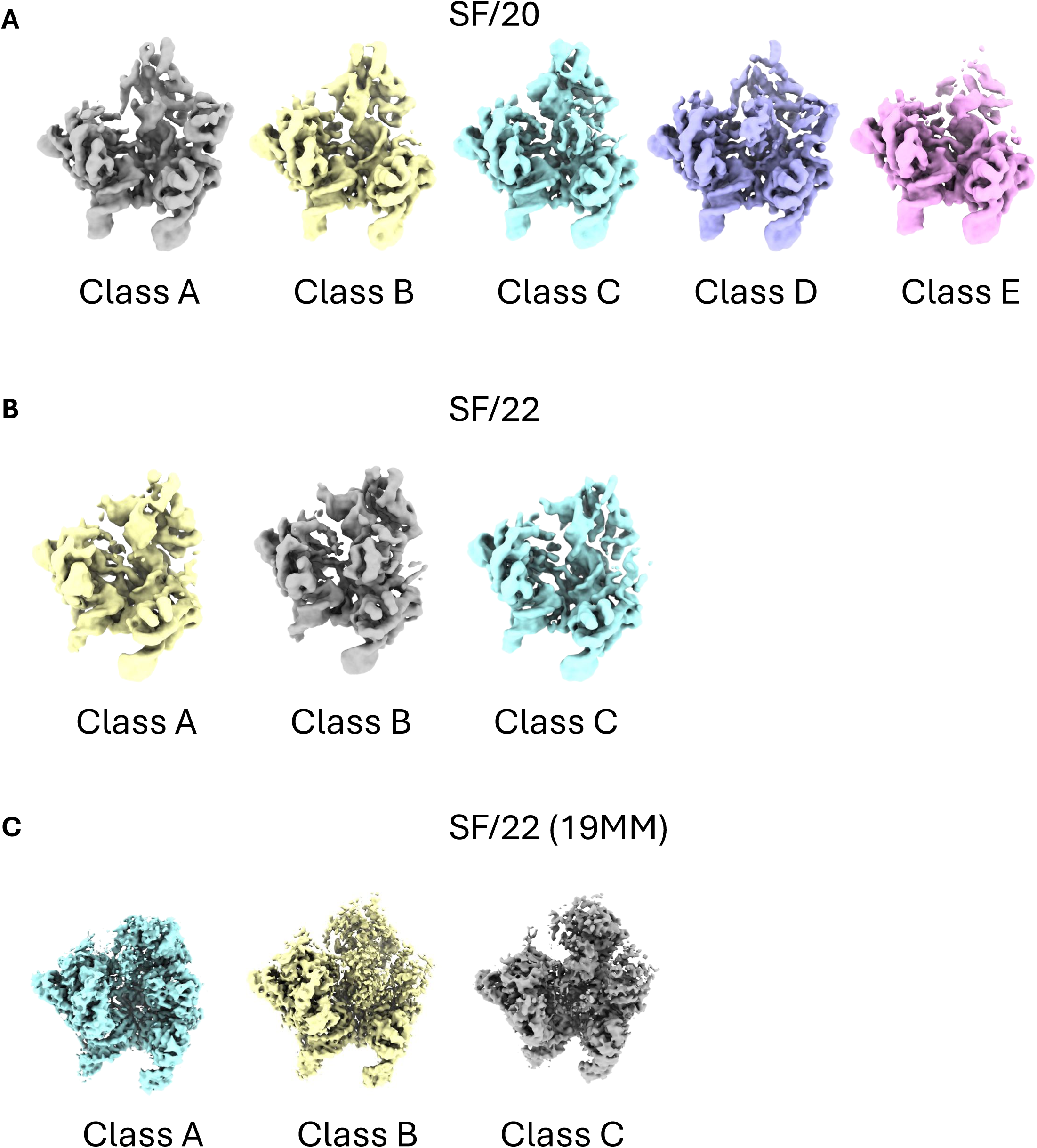

**sFig. 17.**
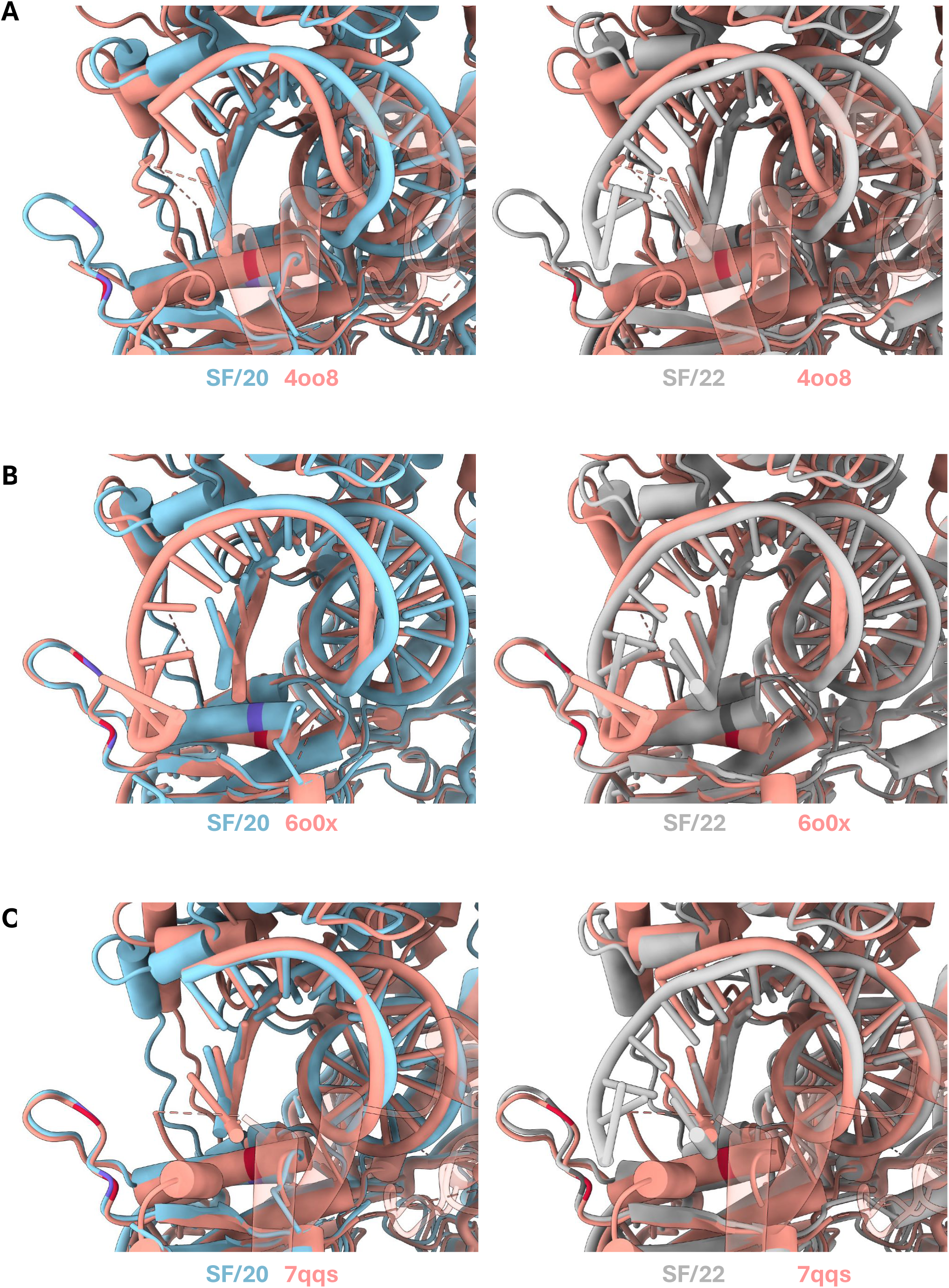

**sFig. 18.**
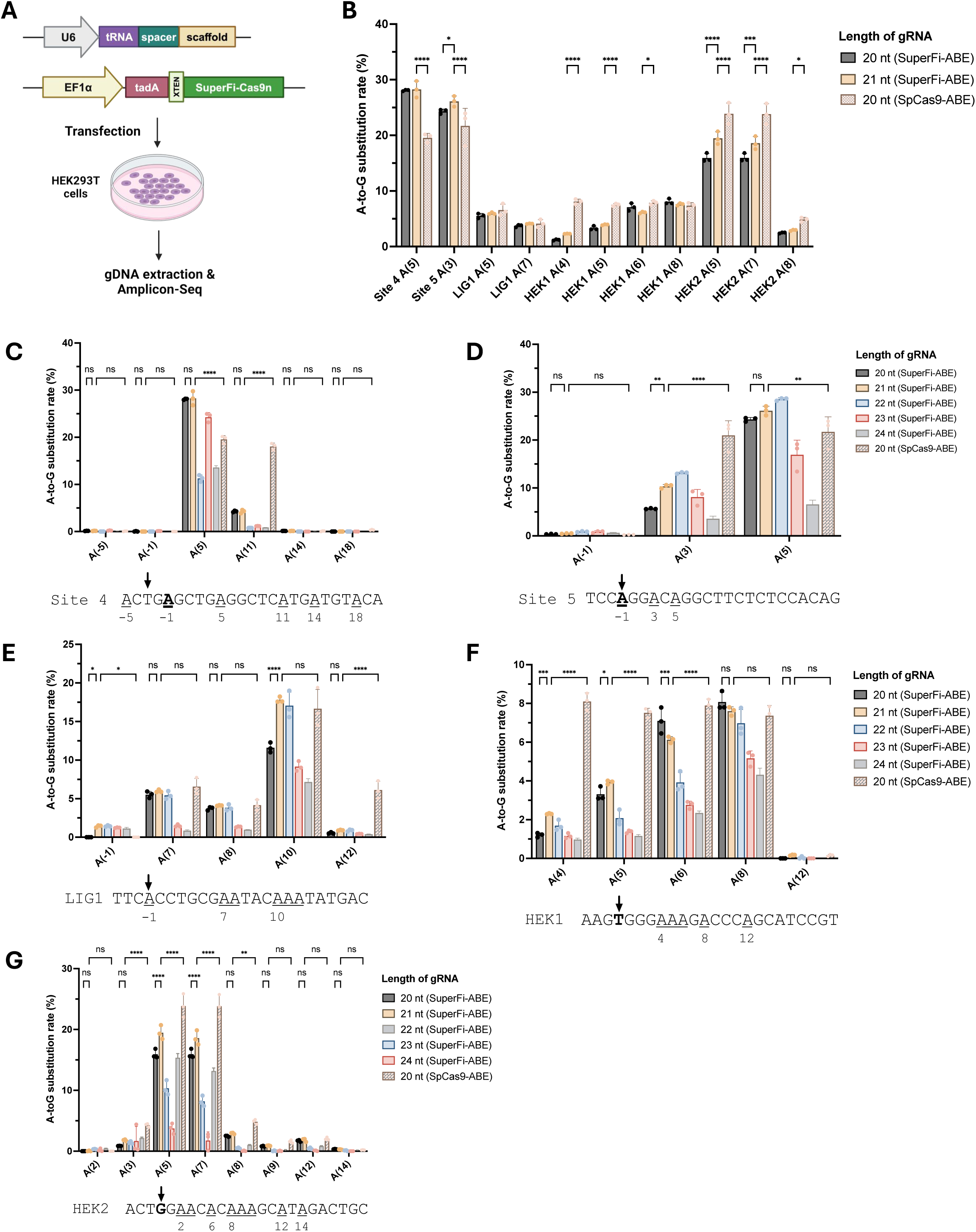

**sFig. 19.**
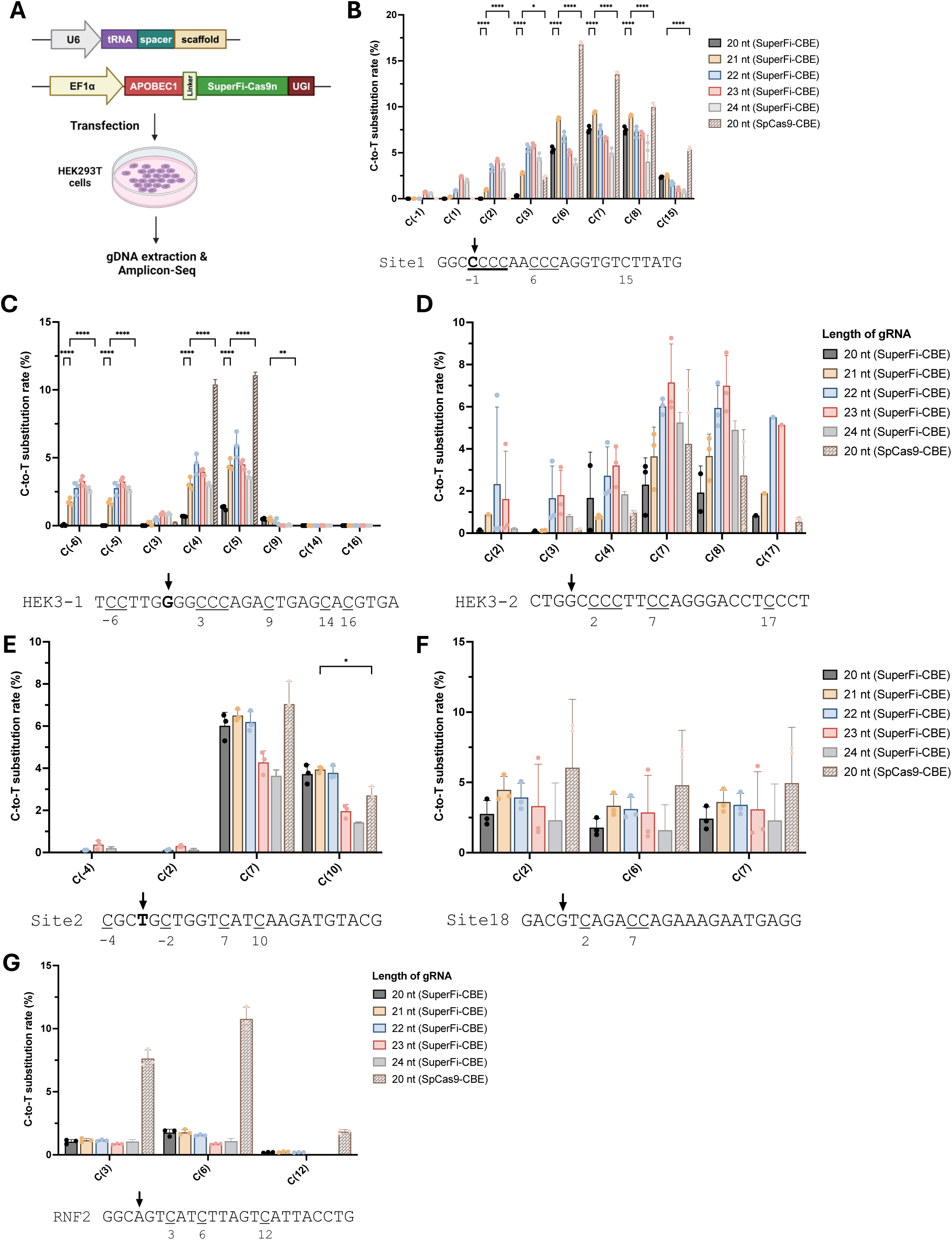

**sFig. 20.**
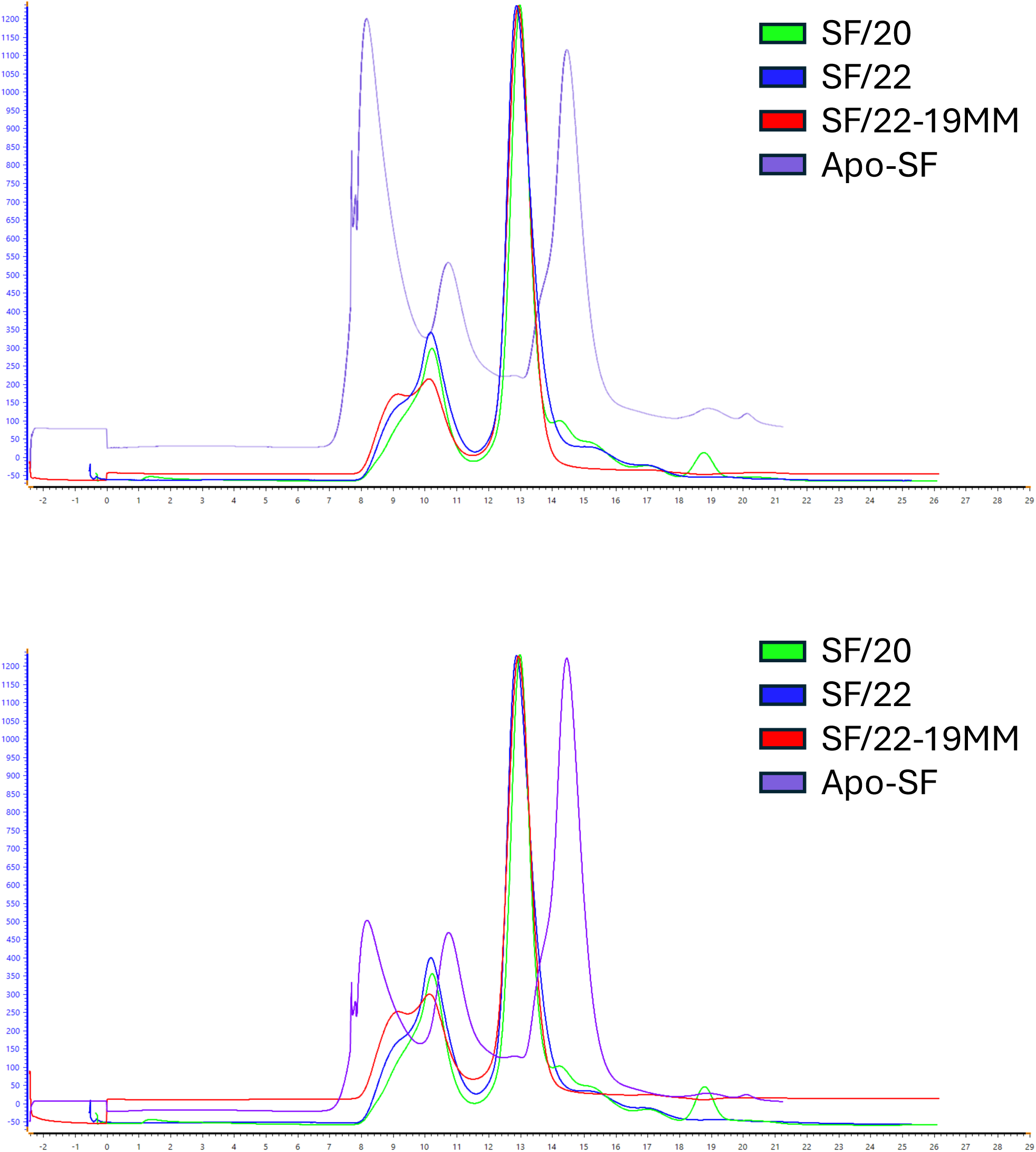

