## Supplementary material for "Spacer Extension Reconciles Specificity and Activity in High-Fidelity Cas9 Genome Editing": Figure legend

**Fig. 1**

A. An experimental scheme of high-throughput profiling on editing efficiencies. SgRNA and the corresponding targeting sequences, including the upstream and downstream flanking sequences, were cloned into a lentivirus backbone and then integrated into the genome of K562 cells by lentivirus transduction. G: The guanine required by the U6 promoter; TS: target sequence; up: upstream 20 bp in the genome; down: downstream 20 bp in the genome.

B. Along the x-axis are the positions of the non-target strand. The nucleotide preferences of SuperFi-Cas9 (up) and SpCas9 (down) in the 63-bp target area (including the flanking sequences) were plotted at the y-axis. At each position along the x-axis, we ranked the target sequence in descending order according to the indel frequencies. The percentage of a particular nucleotide in the first quartile at each position was divided by its percentage in the fourth quartile. The log-2 transformed relative ratio is denoted as the log-odds score (y-axis), representing the relative abundance of that nucleotide in the 1$^{st}$ and 4$^{th}$ quartiles.

C. A scheme shows the complementary between sgRNA and the target sequence.

D. The relative indel frequencies of sgRNA-target pairs in the SuperFi-Cas9 library. The indel frequencies of the top 2,000 sgRNA were transformed into relative indel infrequencies by fitting the numbers into a zero-to-one distribution.

**Fig. 2**

A. An experimental scheme of using 5′-extended sgRNA to edit ten genomic loci in K562 cells with stable SuperFi-Cas9 expression. A tRNA between the U6 promoter and spacer allows the generation of sgRNA with desired lengths.

B. The indel frequencies of sgRNAs with different spacer lengths at ten genomic loci in K562 cells. The sgRNAs were grouped by loci.

C. The indel frequencies of sgRNAs with different spacer lengths at ten genomic loci in K562 cells. The sgRNAs were grouped by length.

D. An experimental scheme of using 5'-extended sgRNA to edit ten genomic loci in HEK293 cells. The SuperFi-Cas9 expression cassette was placed downstream of the sgRNA expression cassette. A tRNA between the U6 promoter and spacer allows the generation of sgRNA with desired lengths.

E. The indel frequencies of sgRNAs with different spacer lengths at ten genomic loci in HEK293T cells. The sgRNAs were grouped by loci.

F. The indel frequencies of sgRNAs with different spacer lengths at ten genomic loci in HEK293T cells. The sgRNAs were grouped by length.

G. An experimental scheme of using 5'-extended sgRNA to edit ten genomic loci in K562 cells. SuperFi-Cas9 and sgRNA were complexed as ribonucleoproteins before electroporation.

H. The indel frequencies of sgRNAs with different spacer lengths at three genomic loci in K562 cells. The sgRNAs were grouped by loci. Data were presented as mean±SD. P values were calculated by two-way ANOVA analysis. *P≤0.05; **P≤0.01; *** P≤0.001, **** P≤0.0001.

**Fig. 3**

A. The relative indel frequencies of sgRNAs when mismatches occur at Site 5. Single mismatch was tiled all over a 22-nt sgRNA, and double mismatches were positioned at the most 5' nucleotides of a 22-nt sgRNA and a 24-nt sgRNA.

B. The relative indel frequencies of sgRNAs when mismatches occur at Site 9. Single mismatch was tiled all over a 22-nt sgRNA, and double mismatches were positioned at the most 5' nucleotides of a 22-nt sgRNA and a 23-nt sgRNA.

C. The number of off-target edits on ten endogenous sites detected by GUIDE-seq.

D-G. Sequences of off-targeting edits of *TTR*-1 (D), *TTR*-2 (E), *PCSK9*-1 (F), and *PCSK9*-2 (G) detected by GUIDE-seq.

**Fig. 4**

A. An experimental scheme of high-throughput profiling on editing efficiencies. A tRNA between the U6 promoter and spacer allows the generation of sgRNA with desired lengths.

B. The indel frequency of sgRNAs from the L2 library against different spacer lengths. Targets whose SpCas9 indel frequencies over 50% were included in this analysis. The column "best" plotted the indel frequencies of the best sgRNA on each target.

C. Along the x-axis are the positions of the non-target strand. The nucleotide preferences of SuperFi-Cas9/20-sgRNA, SuperFi-Cas9/21-sgRNA, and SuperFi-Cas9/22-sgRNA in the 63-bp target area (including the flanking sequences) were plotted at the y-axis. At each position along the x-axis, we ranked the target sequence in descending order according to the indel frequencies. The percentage of a particular nucleotide in the first quartile at each position was divided by its percentage in the fourth quartile. The log-2 transformed relative ratio is denoted as the log-odds score (y-axis), representing the relative abundance of that nucleotide in the 1$^{st}$ and 4$^{th}$ quartiles.

D. The indel frequency of sgRNAs from the L2 and L3 libraries against different spacer lengths.

E. Model architecture of AIdit-SuperFi.

F. Comparison of Pearson coefficients between AIdit-SuperFi and baseline models on the test set.

G. AIdit-SuperFi model predicts the indel frequencies of sgRNAs on human protein-coding genes. The numbers of exon-targeting sgRNA (y-axis) above different thresholds of indel frequencies (x-axis) were plotted. The sgRNAs from "the best length" group are sgRNAs with an extended spacer of 21-nt, 22-nt, 23-nt, or 24-nt.

**Fig. 5**

A. Cartoon overview of SuperFi-Cas9 Cryo-EM structures. The full maps were shown for SF/20 and SF/22 according to their Cryo-EM data. No full map was obtained from the SF/22 (19 MM) Cryo-EM data, and class A was representatively shown here.

B. Detailed view of the aligned structures of SuperFi-Cas9/20-sgRNA (SF/20, skyblue), SuperFi-Cas9/22-sgRNA (SF/22, silver), and SpCas9/20-sgRNA (4un3, salmon).

C-D. Molecular surfaces of SF/22 (C) and SF/20 (D) colored by Coulombic Electrostatic Potential. The resulting potential is in units of kcal/(mol·$e^{-1}$) at 298 K. Positive potential regions are in blue, negative potential regions are in red, and natural or low potential regions are in white. The residues K929, K948, and R951 were highlighted in the corresponding darkened color when compared with 4un3.

E. Indel frequencies of SuperFi-Cas9 and their mutants. 20-sgRNA and 22-sgRNA were used to measure the cleavage activity of each nuclease in HEK293T cells.

F. Indel frequencies of SpCas9 and their mutants. 20-sgRNA and 22-sgRNA were used to measure the cleavage activity of each nuclease in HEK293T cells.

**sFig. 1**

A. The density plot of indel frequencies of sgRNAs in the sgRNA-target paired library. Data from SuperFi-Cas9 and the wild-type SpCas9 were shown.

B. Along the x-axis are the positions of the non-target strand. The nucleotide preferences of Sniper-Cas9, HiFi-Cas9, LZ3-Cas9, and SpCas9-NG in the 63-bp target area (including the flanking sequences) were plotted at the y-axis. At each position along the x-axis, we ranked the target sequence in descending order according to the indel frequencies. The percentage of a particular nucleotide in the first quartile at each position was divided by its percentage in the fourth quartile. The log-2 transformed relative ratio is denoted as the log-odds score (y-axis), representing the relative abundance of that nucleotide in the $1^{st}$ and $4^{th}$ quartiles.

C. The relative indel frequencies of sgRNA-target pairs in the SpCas9 library. The indel frequencies of the top 2,000 sgRNA were transformed into relative indel infrequencies by fitting the numbers into a zero-to-one distribution.

**sFig. 2**

The nomenclature of sgRNAs with different spacer lengths and the denoted positions used in this study.

**sFig. 3**

Sequences of off-targeting edits of *SERPINA1* (A), *KLKB1* (B), *IDS* (C), *WAS* (D), HBB (E), and CCR5 (F) detected by GUIDE-seq.

**sFig. 4**

Along the x-axis are the positions of the non-target strand. The nucleotide preferences of SuperFi-Cas9/19-sgRNA, SuperFi-Cas9/23-sgRNA, and SuperFi-Cas9/24-sgRNA in the 63-bp target area (including the flanking sequences) were plotted at the y-axis. At each position along the x-axis, we ranked the target sequence in descending order according to the indel frequencies. The percentage of a particular nucleotide in the first quartile at each position was divided by its percentage in the fourth quartile. The log-2 transformed relative ratio is denoted as the log-odds score (y-axis), representing the relative abundance of that nucleotide in the 1st and 4th quartiles.

**sFig. 5**

Along the x-axis are the positions of the non-target strand. The nucleotide preferences of SuperFi-Cas9 with sgRNA in different lengths in the 63-bp target area (including the flanking sequences) were plotted at the y-axis. Only sgRNAs with a 5′ leading guanine or adenosine were included in the SuperFi-Cas9 Library 3 profiling library (L3). At each position along the x-axis, we ranked the target sequence in descending order according to the indel frequencies. The percentage of a particular nucleotide in the first quartile at each position was divided by its percentage in the fourth quartile. The log-2 transformed relative ratio is denoted as the log-odds score (y-axis), representing the relative abundance of that nucleotide in the 1st and 4th quartiles.

**sFig. 6**

Indel frequencies were plotted against the number of GC pairs at the PAM-distal region for SuperFi-Cas9 (A) and SpCas9 (B). The number at the x-axis was a sum of GC pairs from the sgRNA position 11 to the most 5′ end.

**sFig. 7**

Indel frequencies were plotted against the number of GC pairs at the PAM-distal region for SuperFi-Cas9 when using different lengths of sgRNAs. The number at the x-axis was a sum of GC pairs from the sgRNA position 11 to the most 5 ’end.

**sFig. 8**

Indel frequencies were plotted against the free energy of the sgRNA-TS duplex at the PAM-distal region for SuperFi-Cas9 (A) and SpCas9 (B). The number at the x-axis was a sum of free energy from the sgRNA position 11 to the most 5 ’end.

**sFig. 9**

Indel frequencies were plotted against the free energy of the sgRNA-TS duplex at the PAM-distal region for SuperFi-Cas9 when using different lengths of sgRNAs. The number at the x-axis was a sum of free energy from the sgRNA position 11 to the most 5 'end.

**sFig. 10**

A. Transformer encoder block of AIdit-SuperFi, replace absolute position encoding with KERPLE, replace ReLU with GEGLU.

B. Pretraining of AIdit-SuperFi with the MLM objective.

C. Architecture of deep learning baseline.

D. Architecture of conventional algorithms baselines.

E. Detailed architecture of CNN.

F. Detailed architecture of RNN.

G. Detailed architecture of MLP.

**sFig. 11**

Comparison of Pearson coefficients between AIdit-SuperFi and baseline models on off-target test set, divided by matching type.

**sFig. 12: SuperFi-Cas9/20nt-sgRNA/dsDNA complex**

A. and B. Representative micrograph, 2D classes and process of particle sorting with binned 3 images. The particles red encircled indicate the Cas9/20nt-sgRNA/dsDNA complex.

C. Data collection yielded 6,171 micrographs, from which 2,027,773 particles were auto-picked. Initial 2D classification in RELION sorted 1,393,256 particles into six classes. Among six classes, the well-resolved class (marked as *1) and classes exhibiting preferred orientation (marked as *2) were selected, and these were then independently subjected to further processing. To address preferred orientation issue, the well-resolved class (*1) underwent further 2D classification to isolate classes having less particles (tilted view). These were combined with particles from the preferred orientation class (*2) for independent 3D classifications. Final classes were curated to eliminate redundant conformations, particularly between derivatives of well-resolved class in first round 3D classification (*1). Since 34,828-particle containing map (marked as *3) is overlapped with a highest resolution map (full map, 3.5Å), which was obtained from combined particles in 3D

classification without alignment derived from '*1 '(bottom left), a), '*3 'map is excluded. The final dataset comprises six maps: one full map and five distinct conformational states (classes A-E).
D. Fourier shell correlation (FSC) curves. The resolutions of full maps were estimated based on the FSC=0.143 criterion.
E. Local resolution estimation map.

**sFig. 13: SuperFi-Cas9/22nt-sgRNA/dsDNA complex**

A and B. Representative micrograph, 2D classes and process of particle sorting with binned 3 images. The particles red encircled indicate Cas9/22nt-sgRNA/dsDNA complex.
C. Data collection yielded 3,308 micrographs from the former dataset and 788 micrographs from the later dataset. After image sorting, 3,018 and 732 micrographs were selected, respectively. Auto-picking resulted in 1,357,650 and 219,600 particles from each dataset. Initial 2D classification in RELION yielded two particle sets. Both datasets underwent three rounds of 3D classification. The selected particles from both datasets were merged into a single dataset for ab-initio reconstruction in cryoSPARC with K=3. The best class underwent non-uniform refinement and local refinement, yielding a full map at 3.6 Å resolution (Full map). To analyze the conformational heterogeneity of the full map, heterogeneous refinement (K=3) was performed, resulting in three distinct classes. Each class was independently processed with non-uniform refinement followed by local refinement to achieve the final reconstructions at resolutions of 4.2 Å, 3.9 Å, and 4.0 Å for Class A, B, and C, respectively.
D. Fourier shell correlation (FSC) curves. The resolutions of full maps were estimated based on the FSC=0.143 criterion.
E. Local resolution estimation map.

**sFig. 14: SuperFi-Cas9/22nt-sgRNA_19mm/dsDNA complex**

A and B. Representative micrograph, 2D classes and process of particle sorting with binned 3 images. The particles red encircled indicate Cas9/22nt-sgRNA_19mm/dsDNA complex.
C. Data collection yielded 1,066 micrographs from Set1 and 2,672 micrographs from Set2. Auto-picking resulted in 479,700 and 1,202,400 particles from each dataset, respectively. Initial 2D classification in RELION yielded 381,211 particles from Set1 and 1,010,967 particles from Set2. Both datasets underwent independent 3D classification in RELION. The selected particles from both datasets were merged into a single dataset of 855,206 particles. This combined dataset

underwent two additional rounds of 3D classification, resulting in 694,155 particles for final processing. To analyze the conformational heterogeneity of the dataset, heterogeneous refinement was performed in cryoSPARC (K=3), yielding three distinct classes. Each class underwent sequential non-uniform refinement, CTF refinement, and local refinement to achieve the final reconstructions at resolutions of 3.3 Å, 3.3 Å, and 3.4 Å for Class A, B, and C, respectively.

D. Fourier shell correlation (FSC) curves. The resolutions of full maps were estimated based on the FSC=0.143 criterion.

E. Local resolution estimation map.

**sFig. 15**

Cartoon view of SF/22 (19-MM) Cryo-EM structure. The class B and class C were shown here.

**sFig. 16**

Density maps of subclasses of SF/20 (A), SF/22 (B), and SF/22 (19 MM) (C).

**sFig. 17**

Detailed view of the aligned structures of SuperFi-Cas9 and published SpCas9 (A: 4oo8, B: 6o0x, C: 7qqs). 4oo8 uses a 20-nt sgRNA. 6o0x uses a 20-nt sgRNA with additional 5'-GG ($GGN_{20}$). 7qqs uses a 20-nt sgRNA with additional 5'-GG ($GGN_{20}$) but only one guanine was shown in the structure.

**sFig. 18**

A. An experimental scheme of applying SuperFi-Cas9n and 5′-extended sgRNA in ABE. A plasmid expressing sgRNA and a plasmid encoding TadA8e were co-transfected to HEK293T cells.

B. The A-to-G substitution rates of SuperFi-ABE within the canonical editing window. Data from three genomic loci were shown.

C-G. The A-to-G substitution rates of SuperFi-ABE were tested at Site 4 (C), Site 5 (D), LIG1 (E), HEK1 (F), and HEK2 (G).

**sFig. 19**

A. An experimental scheme of applying SuperFi-Cas9n and 5'-extended sgRNA in CBE. A plasmid expressing sgRNA and a plasmid encoding APOBEC1, SuperFi-Cas9n, and UGI were co-transfected to HEK293T cells.

B-G. The C-to-G substitution rates of SuperFi-CBE were tested at Site1 (B), HEK3-1 (C), HEK3-2 (D), Site2 (E), Site18 (F), and RNF2 (G).

**sFig. 20**

FPLC profile of apo-SuperFi-Cas9 and substrate-bound SuperFi-Cas9 under 260nm and 280nm.
